# Molecular and functional architecture of striatal dopamine release sites

**DOI:** 10.1101/2020.11.25.398255

**Authors:** Aditi Banerjee, Cordelia Imig, Karthik Balakrishnan, Lauren Kershberg, Noa Lipstein, Riikka-Liisa Uronen, Jiexin Wang, Xintong Cai, Fritz Benseler, Jeong Seop Rhee, Benjamin H. Cooper, Changliang Liu, Sonja M. Wojcik, Nils Brose, Pascal S. Kaeser

## Abstract

Dopamine controls striatal circuit function, but its transmission mechanisms are not well understood. We recently showed that dopamine secretion requires RIM, suggesting that it occurs at active zone-like sites similar to conventional synapses. Here, we establish using a systematic conditional gene knockout approach that Munc13 and Liprin-α, active zone proteins for vesicle priming and release site organization, are important for dopamine secretion. Correspondingly, RIM zinc finger and C_2_B domains, which bind to Munc13 and Liprin-α, respectively, are needed to restore dopamine release in RIM knockout mice. In contrast, and different from conventional synapses, the active zone scaffolds RIM-BP and ELKS, and the RIM domains that bind to them, are expendable. Hence, dopamine release necessitates priming and release site scaffolding by RIM, Munc13, and Liprin-α, but other active zone proteins are dispensable. Our work establishes that molecularly simple but efficient release site architecture mediates fast dopamine exocytosis.

## Introduction

Dopamine is a crucial neuromodulator for the control of locomotion, motivation and reward. While there is rich literature on dopamine actions, dopamine signaling mechanisms remain incompletely understood. An important dopamine pathway in the vertebrate brain arises from cell bodies in the ventral midbrain, and their axons prominently project to the striatum. In the striatum, dopamine axons are extensively branched, a single axon covers a large area, and ascending action potentials as well as local regulatory mechanisms are important for dopamine release (Matsuda et al., 2009; Sulzer et al., 2016; Zhou et al., 2001). Striatal dopamine signaling is often considered a volume transmitter with slow and imprecise signaling because the majority of striatal dopamine varicosities lack synaptic specializations, dopamine receptors on target cells are localized away from release sites, and time-scales of the G-protein coupled receptor (GPCR) signaling are orders of magnitude slower than those of ionotropic receptors (Agnati et al., 1995; Descarries et al., 1996; Missale et al., 1998; Uchigashima et al., 2016). Recent studies, however, find that dopamine influences behavior and synapses with sub-second precision (Howe and Dombeck, 2016; Menegas et al., 2018; Yagishita et al., 2014) and dopamine receptor activation can occur rapidly and requires a high dopamine concentration (Beckstead et al., 2004; Courtney and Ford, 2014; Gantz et al., 2018; Marcott et al., 2018), thereby challenging the model of volume transmission.

However, how signaling structures between dopamine releasing and receiving cells are organized to support these precise functions remains largely unknown. We have recently shown that action potential-triggered dopamine release is executed with millisecond precision and requires the presynaptic scaffolding protein RIM (Banerjee et al., 2020; Liu et al., 2018; Robinson et al., 2019). This has led to the working model that dopamine release is mediated at specialized secretory sites, but the organizers of release site structure beyond RIM are not known (Liu and Kaeser, 2019). Alternatively, RIM could operate to mediate dopamine secretion as a soluble protein or through association with vesicles and molecular release site scaffolds may not be important.

At conventional synapses, vesicular exocytosis is ultrafast, triggered by Ca^2+^ and only occurs within the active zone (Kaeser and Regehr, 2014; Südhof, 2012). In addition to RIM, the active zone contains members of five other protein families: Munc13, ELKS, Liprin-α, RIM-BP and Bassoon (Emperador-Melero and Kaeser, 2020; Wang et al., 2016; Wong et al., 2018). Together, these proteins form a molecular machine that is essential for the spatiotemporal control of secretion via three main mechanisms. First, docking and priming, mediated by RIM and Munc13, renders vesicles ready for fast release (Augustin et al., 1999; Betz et al., 2001; Deng et al., 2011; Koushika et al., 2001; Richmond et al., 1999; Siksou et al., 2009; Varoqueaux et al., 2002). Second, the coupling of Ca^2+^ entry to these release-ready vesicles, by strategically positioning Ca^2+^ channels close to these vesicles, is orchestrated by a tripartite complex between RIM-BP, RIM and channels of the Ca_V_2 family (Han et al., 2011; Held et al., 2020; Hibino et al., 2002; Kaeser et al., 2011; Liu et al., 2011; Müller et al., 2012). Third, the active zone orchestrates the organization and function of essential release machinery components including SNARE proteins and lipids, for example phosphatidylinositol 4,5-bisphosphate (PIP_2_) (van den Bogaart et al., 2011; Honigmann et al., 2013; de Jong et al., 2018; Ma et al., 2011; Milosevic et al., 2005; Di Paolo and De Camilli, 2006).

Dopamine release in the striatum is fast and synchronous, and has a high release probability (Banerjee et al., 2020; Liu et al., 2018), suggesting the presence of active zone scaffolds to organize release. Indeed, dopamine axons contain at least some active zone proteins including RIM, ELKS and Bassoon (Daniel et al., 2009; Liu et al., 2018; Silm et al., 2019; Uchigashima et al., 2016). Moreover, removal of RIM specifically from dopamine neurons abolishes evoked dopamine release, while action potential-independent release persists (Liu et al., 2018; Robinson et al., 2019). In contrast, ELKS is dispensable, and roles of other active zone proteins are unknown (Liu and Kaeser, 2019; Liu et al., 2018). Hence, it has remained uncertain whether dopamine axons employ priming and scaffolding mechanisms similar to conventional synapses. Instead, dopamine release may not require the typical complement of active zone machinery, as has been proposed for the release of peptides, catecholamines and other non-synaptic transmitters (Berwin et al., 1998; van de Bospoort et al., 2012; Farina et al., 2015; Held and Kaeser, 2018; van Keimpema et al., 2017; Liu et al., 2010; Renden et al., 2001). Similarly, dopamine release only partially depends on Ca^2+^ entry through Ca_V_2 channels (Brimblecombe et al., 2015), suggesting that mechanisms other than those mediating the specific tethering of Ca_V_2s (Held et al., 2020; Kaeser et al., 2011) are important.

Here, we determined roles of key active zone proteins for dopamine secretion and complement this analysis with assessing roles of RIM domains that interact with these proteins. We find that Munc13 is essential for dopamine release, and rescue establishes that the interplay between RIM and Munc13 primes dopamine-laden vesicles. We further discovered that the scaffolding requirements of dopamine release sites are different from those at classical active zones: ELKS and RIM-BP are entirely dispensable, and the RIM domains that bind to them are not needed for rescue. Notably, Liprin-α2/3 knockout leads to a ∼50% impairment in dopamine release, and loss of dopamine release in RIM knockouts is efficiently restored by re-expressing a fusion construct of the RIM zinc finger (which binds to Munc13) with the RIM C_2_B domain (which binds to Liprin-α). We conclude that dopamine release sites are relatively simple molecular scaffolds that employ Munc13-mediated vesicle priming for fast release and rely on RIM, Munc13 and Liprin-α for release site scaffolding.

## Results

### RIM domains cooperate in dopamine release

RIM is essential for striatal dopamine secretion, but the molecular mechanisms of RIM function in dopamine release are unknown. At conventional synapses, RIM initiates vesicle priming via recruiting and activating Munc13, to which it binds with its N-terminal zinc finger domain (Fig. 1A) (Andrews-Zwilling et al., 2006; Betz et al., 2001; Deng et al., 2011; Kaeser and Regehr, 2017). Its central PDZ domain directly binds to Ca_V_2 channels and tethers them together with RIM-BP, which binds to a short proline-rich region in RIM positioned between the C-terminal C_2_ domains (Hibino et al., 2002; Kaeser et al., 2011). It’s two C_2_ domains, C_2_A and C_2_B, both bind to PIP_2_, and the C_2_B domain also binds to Liprin-α and is essential for RIM function (de Jong et al., 2018; Koushika et al., 2001; Schoch et al., 2002).

**Figure 1.**
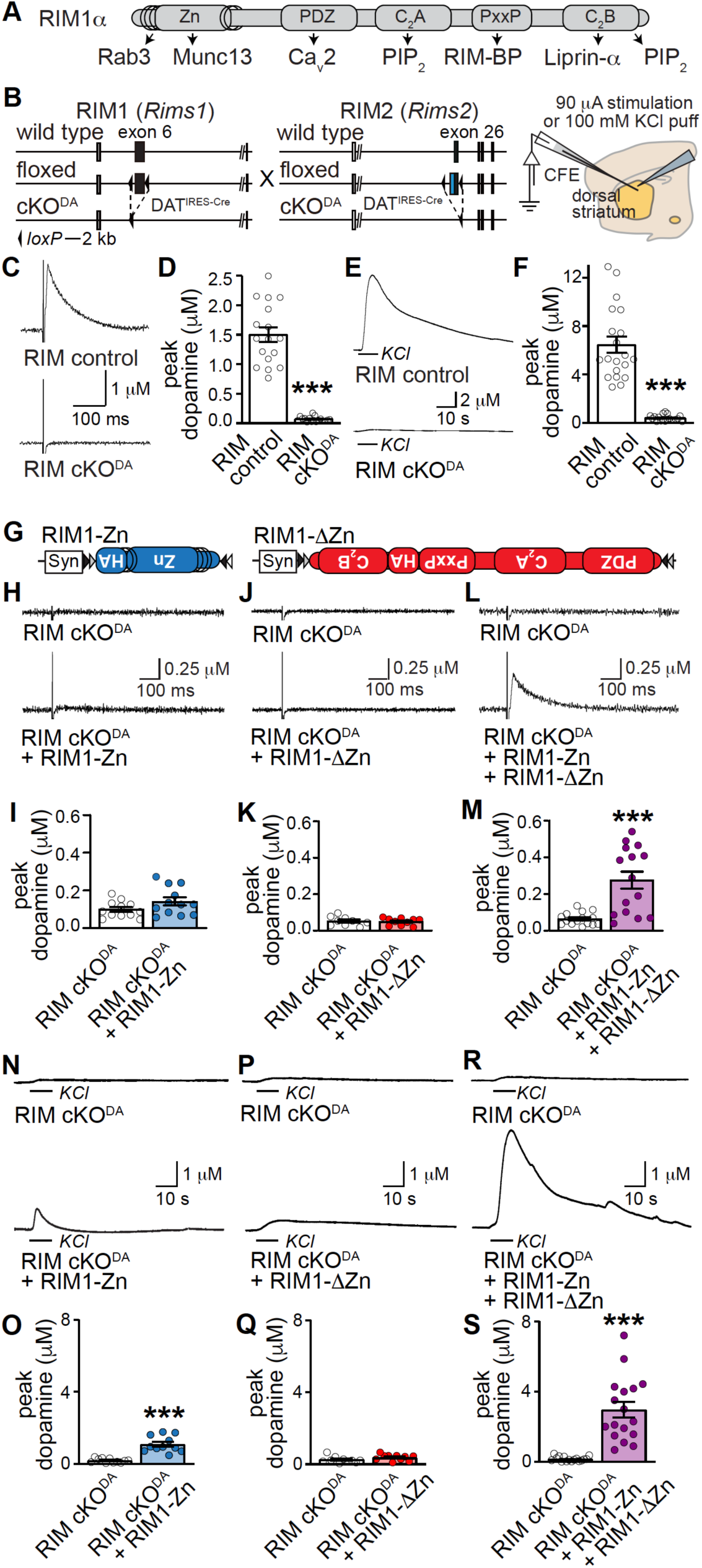
RIM N- and C-terminal domains are necessary for dopamine release **(A)** Schematic of the domain structure of RIM1α and known protein interactions. **(B)** Strategy for ablation of RIM1 and RIM2 in dopamine neurons (RIM cKO^DA^) by crossing floxed RIM1/2 mice to DAT^IRES-cre^ mice (left) and schematic of the slice recording. **(C, D)** Sample traces (C, single sweeps) and quantification (D) of dopamine release evoked by a 90 µA electrical stimulus and measured by carbon fiber amperometry in slices of the dorsolateral striatum, RIM control n = 17 slices/3 mice; RIM cKO^DA^ n = 17/3. **(E, F)** Sample traces (E) and quantification (F) of dopamine release evoked by a local 100 mM KCl puff, RIM control n = 20/3; RIM cKO^DA^ n = 20/3. **(G)** Schematic of AAV5 rescue viruses injected either alone or together into SNc. **(H-L)** Sample traces (H, J and L, single sweeps) and quantification (I, K and M) of dopamine release evoked by a 90 µA electrical stimulus in slices of the dorsolateral striatum, I: RIM cKO^DA^ = 12/4, RIM cKO^DA^ + RIM1-Zn = 12/4; K: RIM cKO^DA^ = 9/3, RIM cKO^DA^ + RIM1-ΔZn = 10/3; M: RIM cKO^DA^ = 15/4, RIM cKO^DA^ + RIM1-Zn + RIM1-ΔZn = 15/4. **(N-R)** Same as H-L, but for a local 100 mM puff of KCl, O: RIM cKO^DA^ = 11/4, RIM cKO^DA^ + RIM1-Zn = 11/4; Q: RIM cKO^DA^ = 10/3, RIM cKO^DA^ + RIM1-ΔZn = 10/3; S: RIM cKO^DA^ = 16/4, RIM cKO^DA^ + + RIM1-Zn + RIM1-ΔZn = 17/4. Data are mean ± SEM, *** p < 0.001 as assessed by Mann Whitney test. For recordings of unrelated wild type mice in each rescue experiment and Ca^2+^ dependence of KCl-triggered release, see Fig. S1.

We started assessing release mechanisms by rescue of RIM knockout phenotypes in striatal slices using amperometric recordings. We confirmed that conditional knockout of RIM in dopamine neurons (RIM cKO^DA^), generated by crossing floxed alleles for both RIM genes to DAT^IRES-cre^ mice (Backman et al., 2006; Kaeser et al., 2008, 2011; Liu et al., 2018), lack action-potential triggered dopamine release (Figs. 1B-1F). Adeno-associated viruses (AAVs) do not allow expression of full-length RIM for rescue as RIM exceeds the viral packaging limit. We instead re-expressed either the RIM zinc finger domain (RIM1-Zn) or the C-terminal scaffolding sequences (RIM1-ΔZn), which together account for the known RIM domains. Rescue expression was restricted to dopamine neurons through the use of cre-dependent AAVs (Fig. 1G) delivered by stereotaxic injection at postnatal days 25 (P25) to P45. Five weeks or more after injection, we assessed dopamine release in RIM cKO^DA^ mice injected either with a control virus, or after re-expression of RIM1-Zn, RIM1-ΔZn, or both. Each experiment was performed such that two RIM cKO^DA^ mice (one with and one without rescue) were recorded on the same day and with the same carbon fiber (Figs. 1H-1S). An unrelated control mouse was also used for recordings on the same day to establish stable detection of dopamine transients (Figs. S1A-S1F).

Expression of RIM1-Zn or RIM1-ΔZn alone showed no rescue of dopamine release evoked by electrical stimulation when compared to RIM cKO^DA^ (Figs. 1H-1K). This is surprising because these constructs are sufficient to restore some action potential-evoked exocytosis of synaptic vesicles or of neuropeptides (Deng et al., 2011; Kaeser et al., 2011; Persoon et al., 2019). However, when RIM1-Zn and RIM1-ΔZn were co-expressed, we observed partial rescue of dopamine release (Figs. 1L-1M). We next tested whether dopamine release in response to depolarization triggered by local puffing of KCl, which depends on the presence of extracellular Ca^2+^ (Figs. S1G and S1H) and is absent in RIM cKO^DA^ mice (Figs. 1E, 1F) (Liu et al., 2018), could be restored. RIM1-Zn alone mediated a small amount of KCl triggered release, and the combined expression led to robust rescue, but RIM1-ΔZn alone was inactive (Figs. 1N-1S). These data establish that the RIM cKO^DA^ phenotype is partially reversible, and both the RIM zinc finger and RIM scaffolding domains are needed for action potential-triggered dopamine release. This is different from regular synapses where each of these RIM fragments mediates partial rescue on its own (Deng et al., 2011; Kaeser et al., 2011).

### Munc13-1 forms small clusters in striatal dopamine axons

We next set out to systematically test the loss-of-function phenotypes of RIM-interacting active zone proteins that bind to the RIM zinc finger or the scaffolding domains. The zinc finger may enhance fusogenicity of dopamine-laden vesicles through vesicle priming. At synapses, vesicle priming is executed by Munc13, which is recruited and activated by RIM zinc finger domains (Andrews-Zwilling et al., 2006; Augustin et al., 1999; Betz et al., 1998; Deng et al., 2011). Given the prominent role of RIM in dopamine release, we hypothesized that dopamine vesicles are primed by Munc13.

Of the three major Munc13 isoforms in the brain (Munc13-1, -2 and -3), Munc13-1 is strongly expressed in midbrain dopamine neurons, while the other Munc13s may be present at low levels (Lein et al., 2007; Saunders et al., 2018). Munc13-1 is present in dopaminergic synaptosomes (Liu et al., 2018), but a lack of suitable antibodies has prevented the assessment of Munc13 distribution in intact striatum using superresolution microscopy. To circumvent this caveat, we used mice in which endogenous Munc13-1 is tagged with EYFP (Fig. 2A) (Kalla et al., 2006). We stained striatal brain sections with anti-GFP antibodies and assessed signal distribution in tyrosine hydroxylase (TH)-labeled dopamine axons using three dimensional structured illumination microscopy (3D-SIM) (Gustafsson et al., 2008; Liu et al., 2018). As expected for an active zone protein present at most synapses, striatal Munc13-1 distribution is broad and Munc13 is present in small clusters (Figs. 2B-2D). The density and shape of TH signals was similar between Munc13-1-EYFP and controls (Figs. 2E-2F), and we used a 40% volume overlap criterion as established before (Liu et al., 2018) to identify Munc13-1 clusters localized within TH axons. There was one Munc13-1 cluster per ∼4 µm of TH axon, and the average volume of each cluster was ∼0.01 µm^3^ (Figs. 2G-2I). We used local shuffling of Munc13-1 objects to asses characteristics of the clusters and signal specificity. Munc13-1 cluster density within dopamine axons dropped after shuffling, establishing that Munc13-1 clusters were more frequently present within TH axons than in their immediate environment. Furthermore, Munc13-1 clusters within TH axons were larger than the clusters detected after shuffling (which represent Munc13-1 clusters of close-by conventional synapses). These findings establish that Munc13-1 is present in active zone-like clusters within dopamine axons.

**Figure 2.**
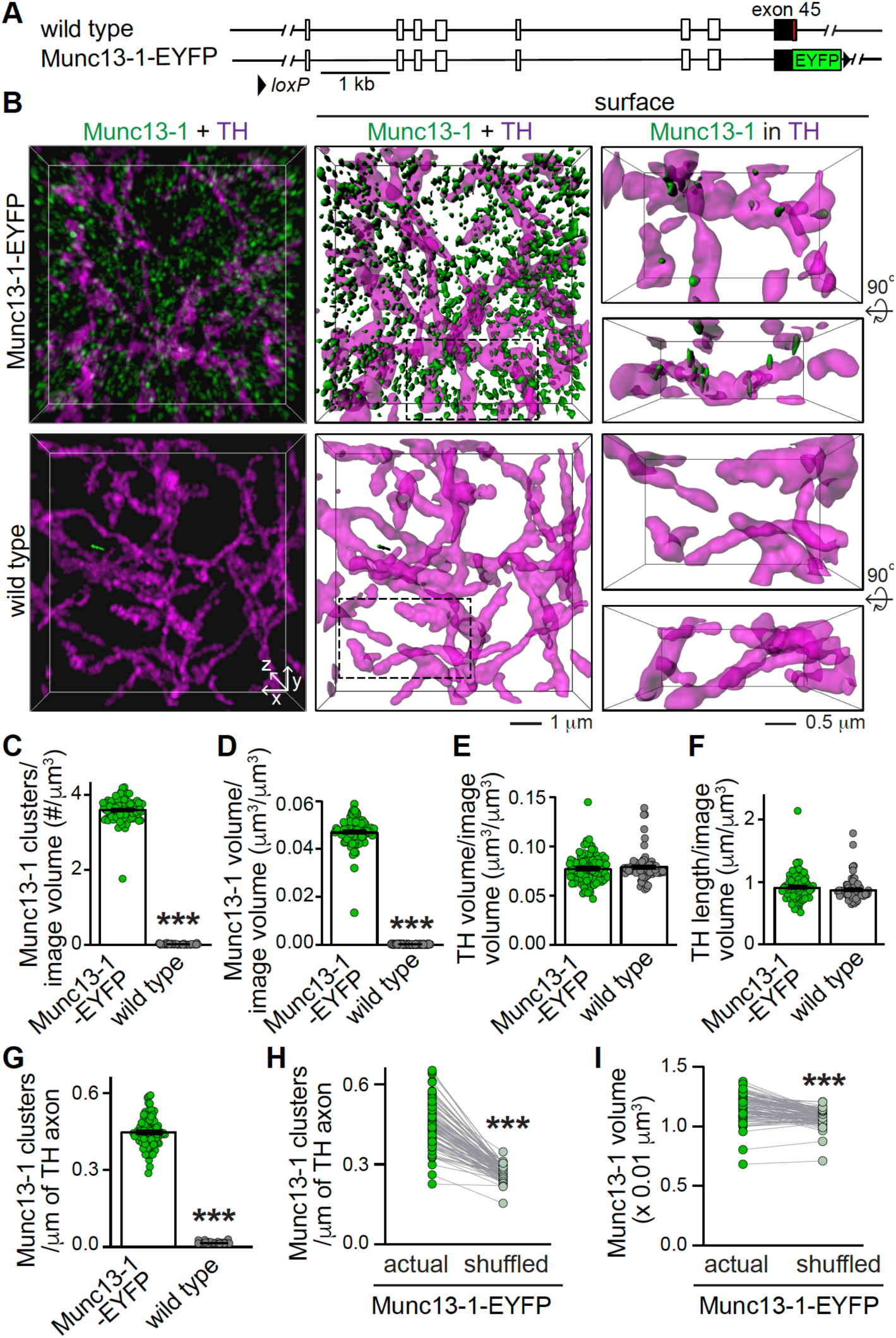
Munc13-1 is expressed in dopamine axons **(A)** Schematic of the Munc13-1-EYFP knock-in mouse (Kalla et al., 2006). **(B)** Sample 3D-SIM images of dorsolateral striatum stained for Munc13-1 and TH in Munc13-1-EYFP and wild type brain sections. GFP antibody staining was used to visualize Munc13-1 and TH was used as a marker of dopamine axons. Volume rendered images (10 x 10 x 2 µm^3^) showing all Munc13-1 and TH (left), surface rendered images of the same volumes (middle) and magnified view (5 x 3 x 2 µm^3^, both frontal view and rotated by +90° along x axis) of only Munc13-1 within TH (right, dotted rectangle in middle, only Munc13-1 with >40% volume overlap with TH are shown). **(C-I)** Quantification of overall number of Munc13-1 clusters per image volume (C), volume occupied by Munc13-1 clusters per image volume (D), TH per image volume (E), length of TH axon per image volume (F), Munc13-1 clusters within TH axons (G), and comparison of Munc13-1 densities (H) and volumes (I) before and after shuffling. For (H) and (I) each Munc13-1 object was randomly shuffled 1000 times within a volume of 1 x 1 x 1 µm^3^, and the actual Munc13-1 densities and volumes were compared to the averaged result after shuffling, Munc13-1-EYFP = 88 images/3 mice; wild type = 86/3. Data are mean ± SEM, *** p < 0.001 as assessed by Mann Whitney test in C, D, E, F, G and Wilcoxon test in H and I.

### Munc13 is essential for evoked striatal dopamine release

To assess roles of Munc13 in dopamine release, we generated new mouse mutants for triple-deletion of Munc13 in vivo. We circumvented lethality of constitutive Munc13-1 deletion through the generation of Munc13-1 conditional knockout mice. We used gene targeting in embryonic stem cells and flanked exon 21 by *loxp* sites (Figs. S2A, S2B). The Munc13-1 floxed mice expressed normal levels of Munc13-1 (Figs. S2C, S2D). We generated constitutive knockout mice by germline-recombination, and full-length Munc13-1 was removed as assessed by Western blotting (Figs. S2E, S2F). A small amount of Munc13-1 (estimated to be less than 5%) at a slightly lower molecular weight persisted and likely lacks exons 21 and 22 (Fig. S2E). In cultured autaptic neurons of these mice, excitatory synaptic transmission was strongly impaired but not abolished (Fig. S3), with a dramatic reduction in the readily releasable pool assessed by sucrose (Figs. S3F, S3G) but a higher initial evoked EPSC amplitude (Figs. S3D, S3E) than previously described for constitutive Munc13-1 knockout mice (Augustin et al., 1999). This difference may be due to the persistence of a small amount of the smaller and possibly partially active Munc13-1 variant in the new mutant (Figs. S2E, S2F).

To assess the necessity for Munc13 proteins in dopamine release, we crossed the Munc13-1 conditional allele to constitutive Munc13-2 and Munc13-3 knockout mice (Augustin et al., 2001; Varoqueaux et al., 2002) and to DAT^IRES-cre^ mice (Backman et al., 2006). We generated Munc13 cKO^DA^ mice (Munc13-1^f/f^ x Munc13-2^-/-^ x Munc13-3^-/-^ x DAT^IRES-cre +/cre^) and Munc13 control mice (Munc13-1^+/f^ x Munc13-2^+/-^ x Munc13-3^-/-^ x DAT^IRES-cre +/cre^) from the same crossings for direct comparison (Fig. 3A). To selectively activate dopamine axons, we expressed a fast version of channelrhodopsin (oChIEF) specifically in dopamine neurons using cre-dependent AAVs delivered by stereotaxic injections (Fig. 3B) at P29-P36 (Banerjee et al., 2020; Lin et al., 2009; Liu et al., 2018). Three or more weeks after injection, we prepared acute brain slices and measured dopamine release triggered by optogenetic activation. In Munc13 control mice, dopamine release appeared normal and very similar in extent to other control mice ((Liu et al., 2018) and Figs. 6, 7). The amplitudes strongly depressed during brief stimulus trains, indicative of a high initial release probability as described before (Liu et al., 2018), and release was abolished by the sodium channel blocker tetrodotoxin (TTX, Fig. 3C) establishing that optogenetic stimulation triggers release via the induction of action potentials. Strikingly, in Munc13 cKO^DA^ mice dopamine release was nearly completely absent, and there was no buildup of release during short stimulus trains (Figs. 3C-3E), although oChIEF-mediated action potential firing was readily elicited in these mice (Fig. S4). To assess release triggered by strong depolarization, we applied a local KCl application (100 mM, 10 s, Figs. 3F-3H). In Munc13 control mice, a strong increase in extracellular dopamine that lasted for tens of seconds and had a ∼3.5-fold larger amplitude than release evoked by optogenetic stimulation was observed. In Munc13 cKO^DA^ mice, no detectable increase in extracellular dopamine was present. Hence, Munc13 is essential for depolarization-induced dopamine release and even strong stimulation fails to elicit significant dopamine release.

**Figure 3.**
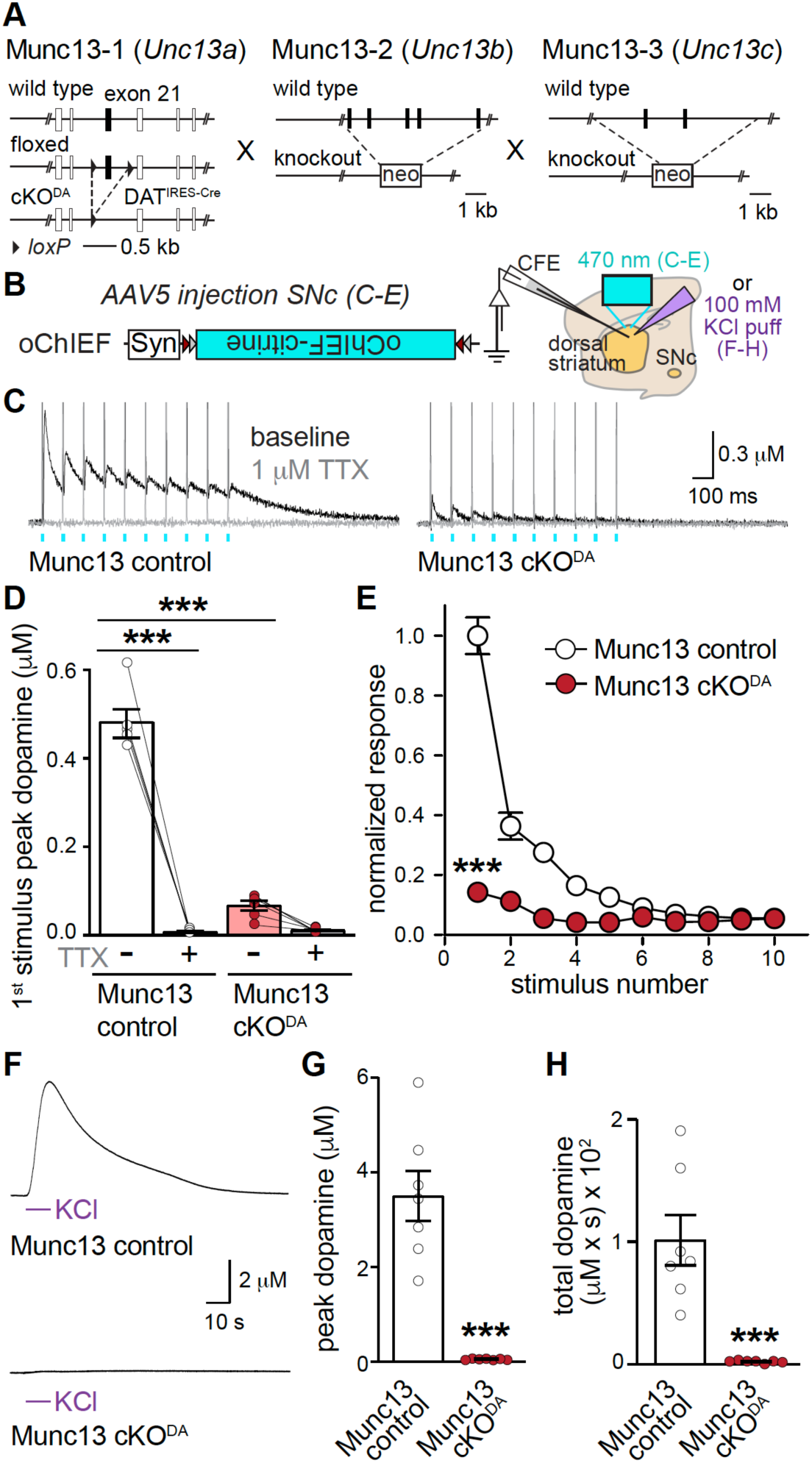
Munc13 is essential for action potential-triggered dopamine release **(A)** Targeting strategy for deletion of Munc13-1, Munc13-2 and Munc13-3 (Munc13 cKO^DA^). Munc13-1 was specifically deleted in dopamine neurons by crossing DAT^IRES-cre^ mice (Backman et al., 2006) to newly generated floxed Munc13-1 mice. Munc13-2 and Munc13-3 constitutive knockout strategies are illustrated as described (Augustin et al., 2001; Varoqueaux et al., 2002). **(B)** Schematic outlining Cre-dependent expression of channelrhodopsin for dopamine neuron activation in Munc13-1 control and cKO^DA^ mice. **(C-E)** Sample traces (C, average of 4 sweeps) of dopamine release evoked by ten 1 ms light pulses at 10 Hz before (black) and after TTX (grey), quantification of peak dopamine amplitudes evoked by the 1^st^ stimulus (D) and peak amplitudes normalized to the average 1^st^ peak of Munc13 control (E), D: Munc13 control = 5 slices/3 mice; Munc13 cKO^DA^ = 6/3 E: Munc13 control = 6/3; Munc13 cKO^DA^ = 6/3. **(F-H)** Sample traces (F), and quantification of peak dopamine amplitudes (G) and area under the curve (H) from the beginning of a KCl puff to 50 seconds. Munc13 control = 7/3; Munc13 cKO^DA^ = 7/3; Data are mean ± SEM, *** p < 0.001 as assessed by ANOVA followed by Sidak’s multiple comparisons test in D; two-way ANOVA (***p<0.001 for genotype, stimulus number and interaction) followed by Sidak’s multiple comparisons test in E (*** p < 0.001 for stimulus 1-4, ** p < 0.01 for stimulus 5), and Mann-Whitney test in G and H. For generation and analysis of synaptic transmission of conditional Munc13-1 cKO mice see Figs. S2 and S3, respectively; for extracellular field recordings of action potential firing in Munc13 cKO^DA^, see Fig. S4.

Striatal cholinergic interneurons can trigger dopamine release through activation of nicotinic acetylcholine receptors on dopamine axons independent of somatic action-potential generation (Cachope et al., 2012; Threlfell et al., 2012; Zhou et al., 2001). This mechanism dominates when electrical field stimulation is used, and as much as 90% of the measured release is mediated by cholinergic stimulation and blocked by nicotinic receptor antagonists (Liu et al., 2018). We measured dopamine release triggered by this cholinergic mechanism using electrical stimulation in slices of Munc13 cKO^DA^ and control mice. Similar to release evoked by optogenetic activation to induce dopamine axon firing (Figs. 3C-3E), electrical stimuli at increasing intensities or in response to 10 stimuli at 10 Hz failed to induced robust release in Munc13 cKO^DA^ mice (Figs. 4A-4D).

**Figure 4.**
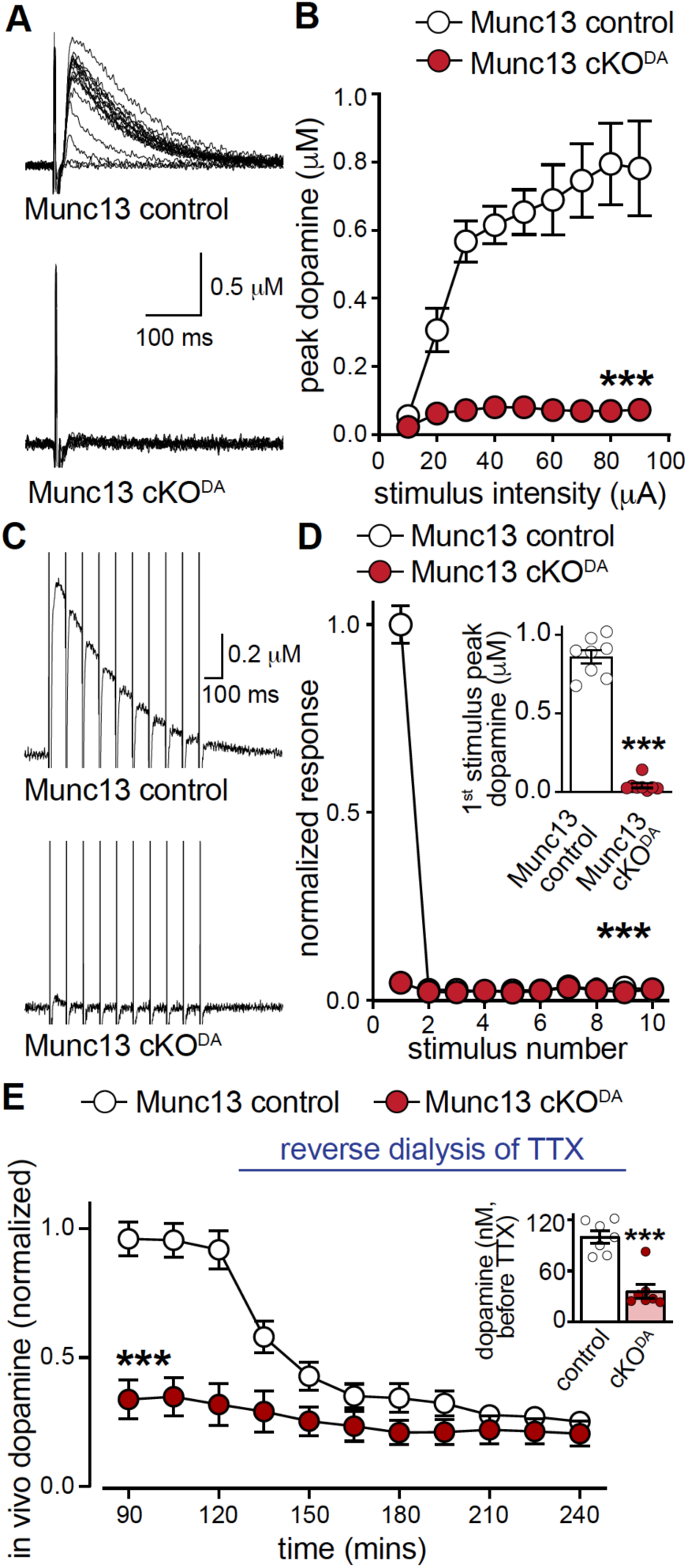
Munc13 deletion abolishes multiple modes of dopamine release **(A, B)** Sample traces of dopamine release (A, single sweeps) and quantification of peak amplitudes (B) evoked by electrical stimulation (10-90 µA single electrical pulses at increasing stimulation intensity). Munc13 control = 10 slices/5 mice; Munc13 cKO^DA^ = 10/5. **(C, D)** Sample traces of dopamine release (C, average of 4 sweeps) and quantification of peak amplitudes normalized to the 1^st^ peak amplitude of Munc13 control (D) in response to ten electrical pulses at 10 Hz train, inset in D shows peak dopamine amplitude for 1^st^ stimulus of the train, Munc13 control = 8/4; Munc13 cKO^DA^ = 8/4. **(E)** Quantification of normalized extracellular dopamine levels within dorsal striatum measured by in vivo microdialysis. All values were normalized to average dopamine values of the 76^th^-120^th^ min of Munc13 control. 10 µM TTX was reverse dialyzed starting at the 121^st^ min, inset shows the mean extracellular dopamine levels from the 76-120^th^ min. Munc13 control and Munc13 cKO^DA^ = 7 mice each. Data are mean ± SEM, *** p < 0.001 as assessed by two-way ANOVA (for genotype, stimulus/time and interaction) in B, D and E followed by Sidak’s multiple comparisons test (B: ** p < 0.01 for 20 µA, *** p < 0.001 for 30-90 µA; D: *** p < 0.001 for 1^st^ stimulus; E: *** p < 0.001 for 90-150^th^ min, ** p < 0.01 for 165-195^th^ min), and unpaired t-tests for insets in D and E.

To determine whether Munc13 is important for dopamine release in vivo, we measured dopamine levels in anesthetized mice using microdialysis (Fig. 4E). In Munc13 control mice, robust extracellular dopamine was detected (Fig. 4E, inset), and reverse dialysis of TTX to inhibit dopamine neuron firing decreased extracellular dopamine to 25% of its initial levels. In Munc13 cKO^DA^, dopamine levels before TTX were strongly reduced. Reverse dialysis of TTX only mildly affected extracellular dopamine, and after TTX the dopamine levels between Munc13 control and cKO^DA^ mice were indistinguishable. We conclude that Munc13 is essential for action potential-triggered dopamine release in vivo. Remarkably, some extracellular dopamine persisted after Munc13 knockout, and this could be due to release that is independent of Munc13 and action potentials similar to RIM-independent release (Liu et al., 2018; Robinson et al., 2019), release mediated by residual Munc13-1 (Fig. S2), or – trivially - tissue damage during microdialysis.

### Roles for Munc13 in dopamine axon structure

A body of literature has established that Munc13 is essential for synaptic vesicle release, but that removal of Munc13 and the resulting block of glutamate release from cultured hippocampal synapses does not impair synaptic structure (Sigler et al., 2017; Varoqueaux et al., 2002). However, it is not known whether this is true for neuromodulatory systems and their development in vivo. We set out to assess whether Munc13 is important for axonal and release site structure in midbrain dopamine neurons. First, we prepared synaptosomes from striatal homogenates of Munc13 control and Munc13 cKO^DA^ mice (Figs. 5A-5E) as we have done before (Liu et al., 2018), which circumvents limitations of quantifying fluorescent signals in tissue densely packed with synapses. We stained synaptosomes with antibodies against the active zone marker Bassoon, the synaptic vesicle protein synaptophysin, and the dopamine neuron marker tyrosine hydroxylase (TH). We generated regions of interest using TH (TH^+^) and synaptophysin (syp^+^) signals, and quantified signal intensities of the various markers within these ROIs. In TH^+^ ROIs of Munc13 cKO^DA^ synaptosomes, TH levels were moderately increased (Fig. 5B), but synaptophysin intensities were somewhat decreased (Fig. 5C), suggesting alterations in their structure. We next assessed Bassoon signals as a proxy for release site scaffolding. Bassoon levels within dopaminergic varicosities (TH^+^/syp^+^ ROIs) were significantly increased (Figs. 5D, 5E), and the same was true in TH^+^ only ROIs in an independent experiment (Figs. S5A, S5B, S5C). Munc13 cKO^DA^ did not affect Bassoon intensities in non-dopamine synapses (TH^-^/syp^+^ ROIs).

**Figure 5.**
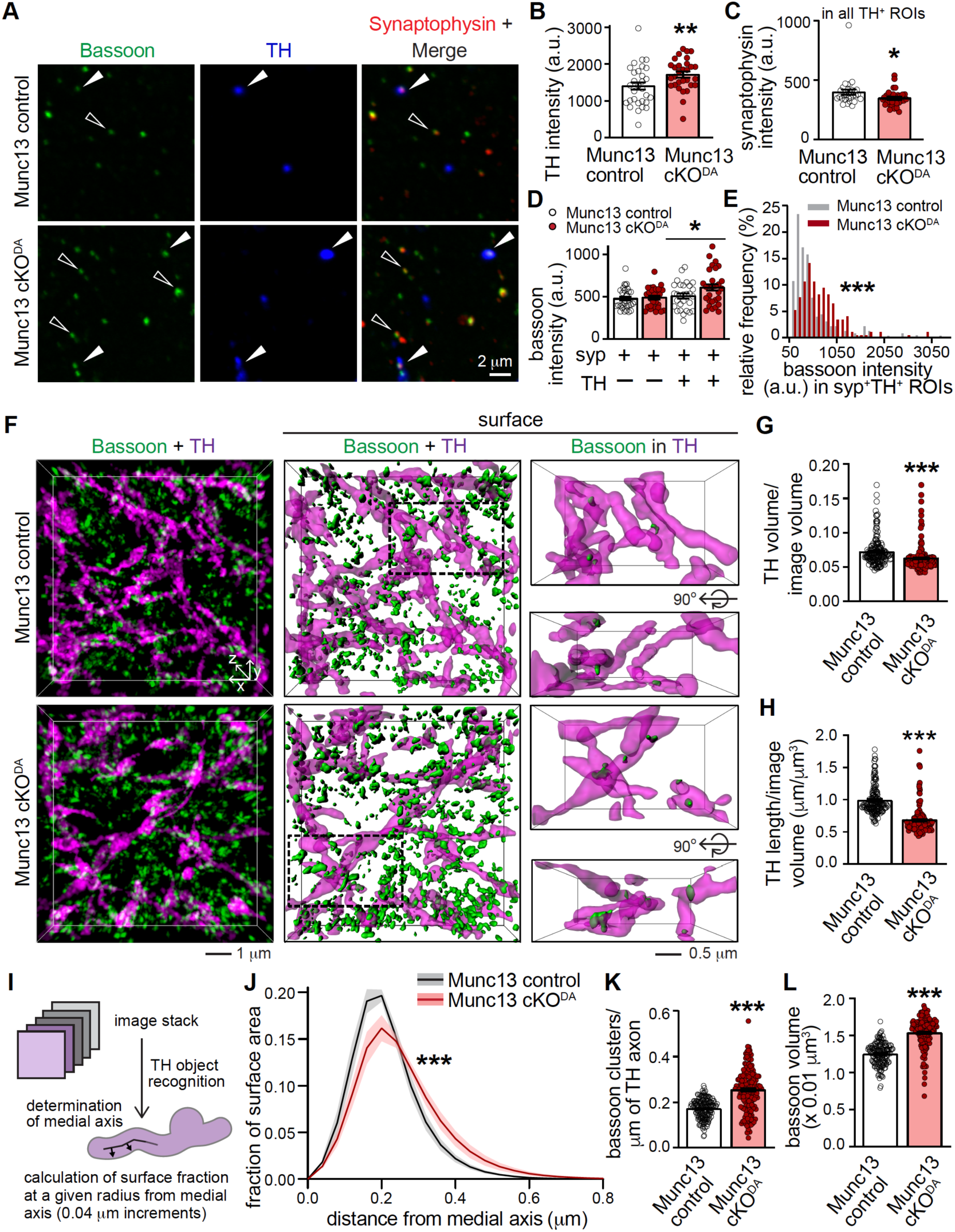
Munc13 ablation in dopamine neurons affects axon structure and bassoon clustering **(A)** Sample confocal images of striatal synaptosomes stained with the active zone marker bassoon, the vesicle marker synaptophysin and TH, with synaptosomes co-expressing all three proteins (solid arrowhead) or non-dopaminergic synaptosomes (hollow arrows, no TH signal) are highlighted. (**B-E**) Quantification of the experiment shown in A, displaying TH intensity in TH^+^ ROIs (B), synaptophysin intensity in TH^+^ ROIs (C), bassoon intensities in ROIs positive for synaptophysin (syp) and/or TH (D), and the frequency distribution histogram for bassoon intensity in syp^+^TH^+^ ROIs (E). Only bassoon intensities within syp^+^TH^+^ ROIs which are greater than 3 times the average intensity of all pixels are shown in D. The frequency histogram in E is plotted for all bassoon intensities within syp^+^TH^+^ ROIs. Munc13 control and Munc13 cKO^DA^ = 30 images /3 mice each. (**F**) Sample 3D-SIM images of dorsolateral striatum stained for bassoon and TH. Volume rendered images (left, 10 x 10 x 2 µm^3^) with all bassoon; surface rendering of the same images with all bassoon (middle) and magnified views (right, 5 x 3 x 2 um^3^, frontal view and rotated by +90° along x axis) of only bassoon within TH (> 40% volume overlap) are shown. **(G, H)** Quantification of the experiment shown in F displaying TH volume per image (G), TH axon length per image volume (H), Munc13 control = 163 images/4 mice; Munc13 cKO^DA^ = 165/4. **(I, J)** Assessment of TH axon shape in the experiment shown in F with outline of the analysis (I) and the proportion of the axonal surface at a specific distance of the medial axis of the TH axon (J), Munc13 control = 160/4; Munc13 cKO^DA^ = 160/4. **(K, L)** Quantification of the density (K) and volume (L) of bassoon clusters (K) localized within TH axons (> 40% volume overlap), n as in G, H. Data are mean ± SEM except for J (mean ± SD), * p < 0.05, ** p < 0.01, *** p < 0.001 as assessed by unpaired t-test in B, C, G, H, K and L; one-way ANOVA followed by Sidak’s multiple comparisons test in D; Kolmogorov-Smirnov test for the data shown in E and Chi square test in J. For quantification of bassoon intensities within TH^+^ ROIs in an independent synaptosome experiment, see Fig. S5.

To assess whether similar changes were present in intact striatum, we used 3D-SIM superresolution microscopy (Figs. 5F-5L). In slices of the dorsolateral striatum, the TH axons appeared less dense but irregular in shape in Munc13 cKO^DA^, the volume occupied by TH was somewhat reduced, and the length of the skeletonized TH axon network per image volume was decreased (Figs. 5G, 5H). When we plotted binned histograms of the radii of TH axons (Figs. 5I, 5J), a right shift in the distribution was detected, indicating that there were more axonal segments with a larger radius likely explaining the increased TH intensities in synaptosomes. We further found that both the densities and volumes of Bassoon clusters were enhanced in Munc13 cKO^DA^ (Figs. 5K, 5L), again matching with the increased bassoon intensities in synaptosomes (Figs. 5D, 5E). We conclude that Munc13 is necessary for normal dopamine axon structure. This role is likely independent of the loss of action potential-triggered dopamine release because ablation of RIM or synaptotagmin-1 in dopamine neurons also abolish evoked release, but do not induce similar axon structural changes (Banerjee et al., 2020; Liu et al., 2018).

### RIM-BP is dispensable for dopamine release

RIM and Munc13 control dopamine release likely via vesicle priming, but this mechanism alone is not sufficient to restore dopamine release in RIM cKO^DA^ neurons as C-terminal RIM sequences are needed (Fig. 1). At conventional synapses, these domains mediate scaffolding, including the tethering of Ca^2+^ channels (Han et al., 2011; Kaeser et al., 2011). RIM contributes to this function in a tripartite complex with Ca_V_2s and RIM-BP (Acuna et al., 2016; Hibino et al., 2002; Kaeser et al., 2011; Liu et al., 2011). It is noteworthy that striatal dopamine release is partially resistant to blockade of Ca_V_2.1 (P/Q-type) and 2.2 (N-type) channels (Brimblecombe et al., 2015), indicating that other Ca^2+^ sources mediate release. RIM-BPs appear to be expressed in dopamine neurons (Lein et al., 2007; Saunders et al., 2018) and would be ideally suited to couple to both Ca_V_2s and other Ca_V_s, for example Ca_V_1s, because the interaction between RIM-BP SH3 domains and proline-rich regions in Ca_V_ C-termini occurs with Ca_V_1s and Ca_V_2s (Hibino et al., 2002). Hence, RIM-BPs may be critical for Ca^2+^ channel organization in dopamine varicosities.

We tested this hypothesis by knockout of RIM-BP1 and RIM-BP2 in dopamine neurons (RIM-BP cKO^DA^, Fig. 6A), crossing “floxed” alleles for these genes (Acuna et al., 2015) to DAT^IRES-cre^ mice (Backman et al., 2006). We measured dopamine release evoked by action potentials via oChIEF (Figs. 6B, 6C), by electrical stimulation which engages cholinergic release triggering (Fig. S6), or KCl depolarization (Figs. 6D-6F). Surprisingly, dopamine release was not affected in RIM-BP cKO^DA^ mice. While we cannot exclude minor roles in the organization of release sites, this establishes that RIM-BPs are largely dispensable for striatal dopamine release, different from hippocampal mossy fiber synapses, the calyx of Held and the fly neuromuscular junction (Acuna et al., 2015; Brockmann et al., 2019; Liu et al., 2011).

**Figure 6.**
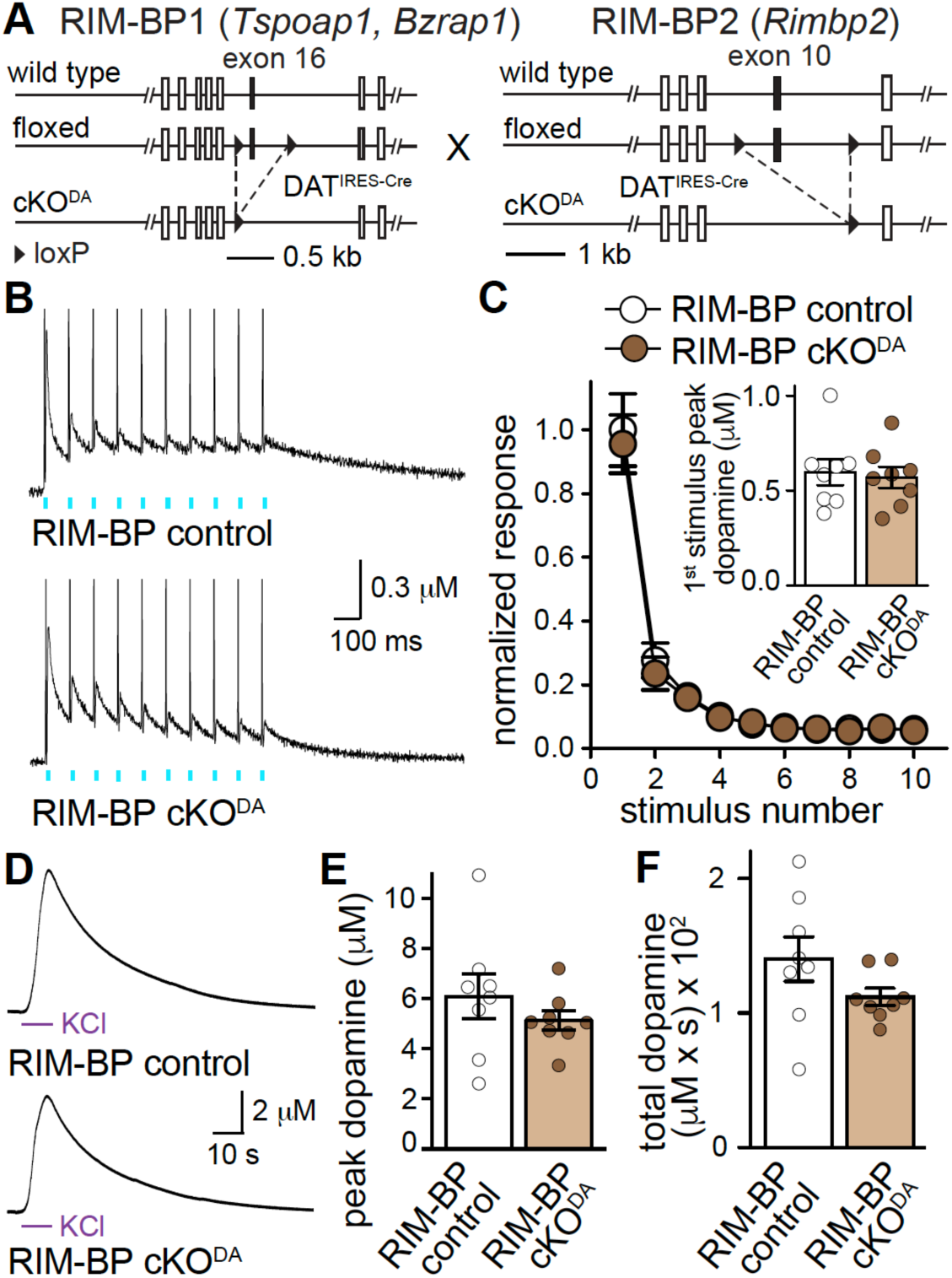
RIM-BP is dispensable for dopamine release **(A)** Strategy for deletion of RIM-BP1 and RIM-BP2 in dopamine neurons (RIM-BP cKO^DA^) using conditional mouse genetics (Acuna et al., 2015; Backman et al., 2006). **(B, C)** Sample traces of dopamine release (B, average of four sweeps) evoked by ten 1 ms- light pulses at 10 Hz and quantification of amplitudes (C) normalized to average of the first peak amplitude of RIM-BP control. Inset in C shows peak amplitude evoked by the first stimulus. RIM-BP control = 8 slices/5 mice; RIM-BP cKO^DA^ = 8/5. **(D-F)** Sample traces (D), quantification of peak amplitudes (E) and area under the curve (F) in response to a KCl puff. RIM-BP control = 8/3; RIM-BP cKO^DA^ = 8/3. Data are mean ± SEM. *** p < 0.001 as assessed by two-way ANOVA (*** for stimulus number) followed by Sidak’s multiple comparisons test in C, unpaired t-test for inset in C, and Mann-Whitney test in E and F. For dopamine release evoked by electrical stimulation in RIM-BP cKO^DA^, see Fig. S6.

### Roles for Liprin-α2 and -α3 in dopamine release

Given that RIM-BP (Fig. 6) and ELKS (Liu et al., 2018) are dispensable for dopamine release, other C-terminal interactions of RIM are likely important. RIM binds to Liprin-α with its C-terminal C_2_B domains (Schoch et al., 2002). Liprin-α proteins are organizers of invertebrate active zones (Böhme et al., 2016; Patel and Shen, 2009; Zhen and Jin, 1999), and we are only beginning to understand vertebrate Liprin-α functions. Vertebrates express four Liprin-α genes (*Ppfia1-4*) that give rise to Liprin-α1 through -α4 proteins. While Liprin-α2 and -α3 are strongly expressed in brain and localized to synapses and active zones, brain Liprin-α1 and - α4 expression is low (Emperador-Melero et al., 2020; Wong et al., 2018; Zürner et al., 2011).

We generated a dopamine-neuron specific double knockout of Liprin-α2 and -α3 (Liprin-α cKO^DA^, Fig. 7A) by crossing floxed Liprin-α2 mice to constitutive Liprin-α3 knockouts and to DAT^IRES-cre^ mice (Backman et al., 2006; Emperador-Melero et al., 2020; Wong et al., 2018). We first measured action potential-triggered dopamine release using optogenetic stimulation, and found that dopamine release was reduced by ∼50% in Liprin-α cKO^DA^ mice (Figs. 7B, 7C). Release triggered by KCl mediated depolarization was reduced similarly (Figs. 7D-7F). In contrast, release triggered by electrical stimulation, which strongly depends on nAChR receptor activation (Liu et al., 2018; Threlfell et al., 2012), appeared unimpaired (Fig. S7). This could be because cholinergic triggering engages different release mechanisms, because of technical differences in the stimulation methods, or because knockout of Liprin-α leads to changes in dopamine-neuron excitability or in cholinergic interneuron function. Altogether, our data suggest important, albeit not essential, roles for Liprin-α in striatal dopamine release, suggesting that C_2_B-mediated scaffolding functions of RIM mediate dopamine secretion.

**Figure 7.**
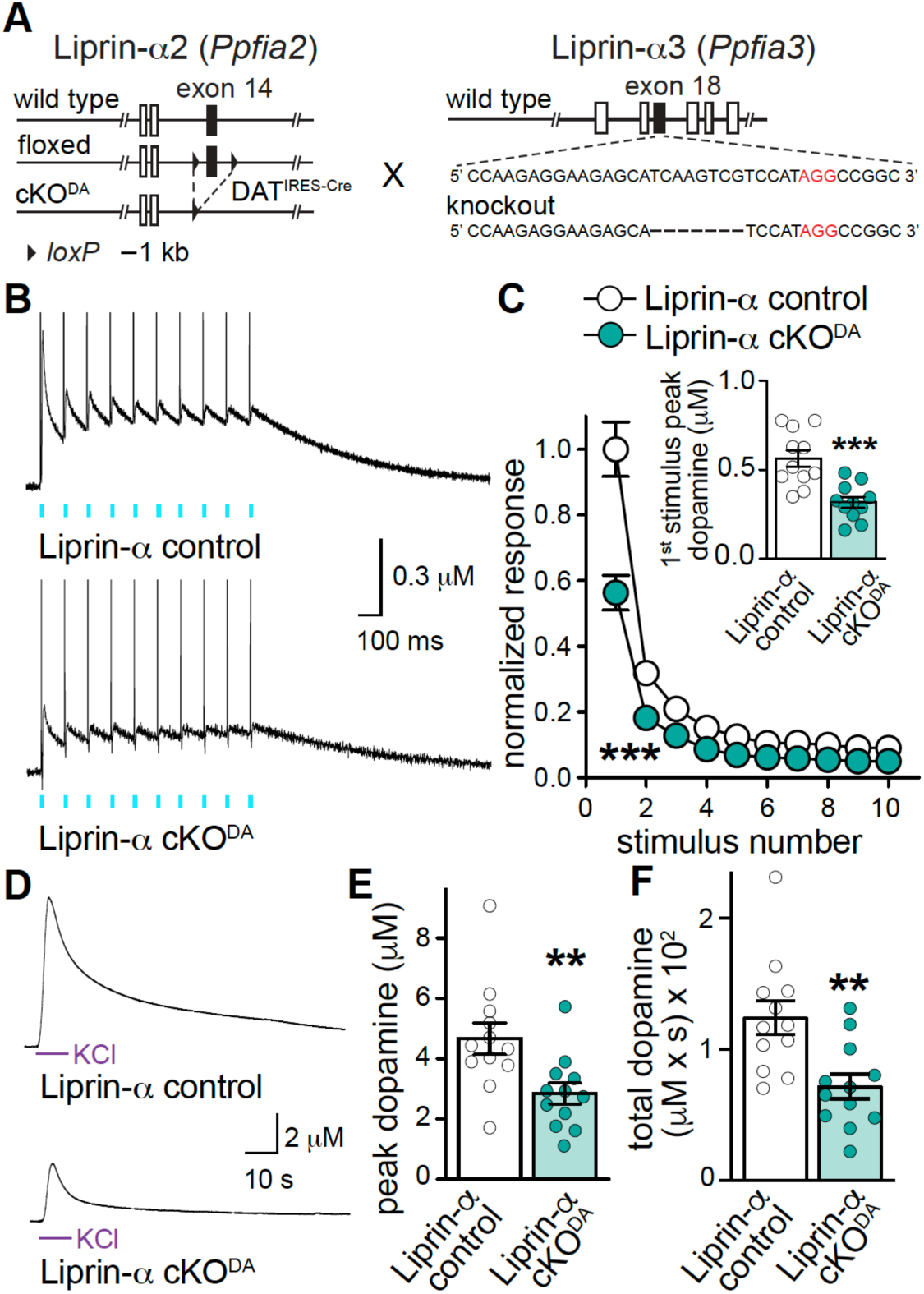
Liprin-α2/3 is important for dopamine release **(A)** Strategy for deletion of Liprin-α2 and Liprin-α3 in dopamine neurons (Liprin-α2/3 cKO^DA^) using mouse genetics (Backman et al., 2006; Emperador-Melero et al., 2020; Wong et al., 2018). **(B, C)** Sample traces of dopamine release (B, average of four sweeps) evoked by ten 1 ms light pulses at 10 Hz and quantification of amplitudes (C) normalized to the average of the first peak amplitude of RIM-BP control. Inset in C shows peak amplitude evoked by the first stimulus. Liprin-α2/3 control = 11 slices/4 mice; Liprin-α2/3 cKO^DA^ = 11/4. **(D-F**) Sample traces (D), quantification of peak amplitudes (E) and area under the curve (F) in response to a KCl puff. Liprin-α2/3 control = 12/8; Liprin-α2/3 cKO^DA^ = 12/8. Data are mean ± SEM. ** p < 0.01, *** p < 0.001 as assessed by two-way ANOVA (*** p < 0.001 for genotype, stimulus number and interaction) followed by Sidak’s multiple comparisons test in C (*** p < 0.001 for first and second stimuli), unpaired t-test for inset in C, and Mann-Whitney test in E and F. For dopamine release evoked by electrical stimulation in Liprin-α2/3 cKO^DA^ see Fig. S7.

### Tethering of dopamine vesicle priming mechanisms to RIM-C_2_B domains restores dopamine release

Our genetic analysis suggests that RIM operates through Munc13 for vesicle priming and through Liprin-α for scaffolding, mediated by the N-terminal zinc finger and C-terminal C_2_B domains, respectively. Interactions of the central RIM domains, including those with ELKS, Ca_V_2s and RIM-BP appear dispensable. If true, dopamine release should be largely restored by the presence of the zinc finger domain to boost fusogenicity via Munc13 (Figs. 1-5) and the RIM C_2_B domain, which may support scaffolding via Liprin-α and PIP_2_ (de Jong et al., 2018; Schoch et al., 2002). We generated a fusion protein of the RIM zinc finger domain with the RIM C_2_B domain, and expressed it using cre-dependent AAVs to test this hypothesis.

Strikingly, fusing the Munc13- and Liprin-α-interacting domains of RIM robustly enhanced dopamine release evoked by electrical stimuli or KCl in RIM cKO^DA^ slices (Figs. 8B, 8C, 8D, 8E). Rescue was as efficient as when all RIM domains were co-expressed for electrically evoked release, but not for KCl-triggered release (Figs. 1, S8C, S8D). These data suggest that RIM-C_2_B domains enable scaffolding of the RIM zinc finger to release sites for action potential-triggered dopamine release. These two domains appear sufficient to mediate the minimally needed release site functions for dopamine, indicating that dopamine release has strikingly limited molecular requirements for release.

**Figure 8.**
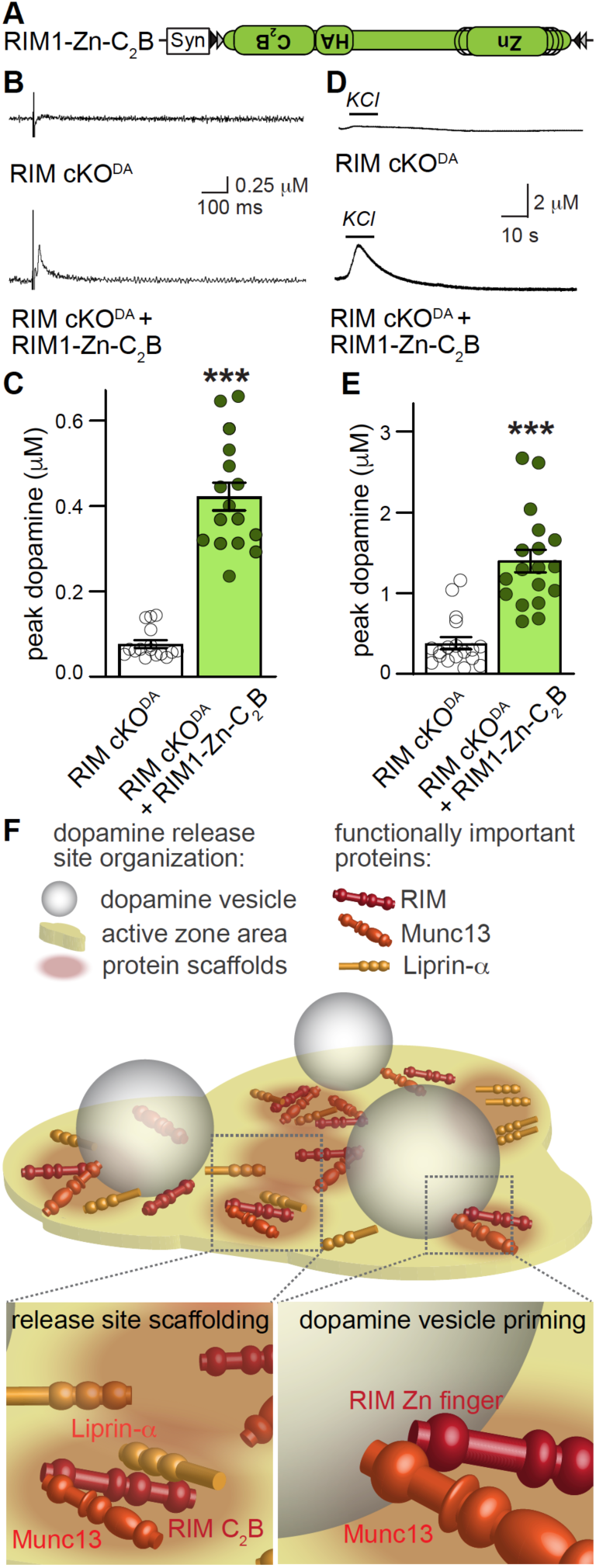
RIM C_2_B domain-mediated scaffolding restores action potential-triggered dopamine release **(A)** Schematic of AAV5 rescue viruses injected into SNc. **(B, C)** Sample traces (B, single sweeps) and quantification of peak dopamine (C) evoked by a 90 µA electrical stimulus in slices of the dorsolateral striatum. RIM cKO^DA^ = 16 slices/4 mice; RIM cKO^DA^ + RIM1-Zn-C_2_B = 16/4. **(D, E)** Sample traces (B) and quantification (C) of peak dopamine evoked by a local 100 mM KCl puff, RIM cKO^DA^ = 18/4; RIM cKO^DA^ + RIM1-Zn-C2B = 18/4. **(F)** Model of active zone-like release sites in dorsal striatum. RIM, Munc13, Liprin-α form release sites in dopamine varicosities, with Munc13 and RIM mediating dopamine vesicle priming and all three proteins contributing to release site scaffolding. Fast, action-potential triggering of dopamine release relies on RIM, Munc13 and the Ca^2+^ sensor synaptotagmin-1 (Banerjee et al., 2020). Notably, only 25-30% of varicosities on dopamine axons have active zone scaffolds (Liu et al., 2018). Data are mean ± SEM. *** p < 0.001 as assessed by Mann-Whitney test in C and E. For recordings of unrelated wild type mice in each rescue experiment see Fig. S8.

## Discussion

Despite central roles for striatal dopamine in circuit regulation and behavior, the molecular and functional organization of its release has remained largely unexplored. Dopamine release requires the active zone protein RIM (Liu et al., 2018; Robinson et al., 2019), but no components or mechanisms of active zone-like release sites are known beyond this requirement. Here, we find that Munc13 is essential for dopamine release and that RIM and Munc13 co-operate to promote dopamine vesicle priming (Figs. 1-5). The scaffolding mechanisms that organize dopamine release sites appear remarkably simple (Figs. 1, 6-8). Most classical active zone scaffolds, including RIM-BP (Fig. 6) and ELKS (Liu et al., 2018), are dispensable for dopamine release. The C-terminal RIM C_2_B domains are important for dopamine release, and may mediate secretion through Liprin-α (Figs. 1, 7, 8). Our data suggest a model (Fig. 8F) in which dopamine release is established through relatively simple active zone like sites: RIM and Munc13 mediate dopamine vesicle priming and operate together with Liprin-α as essential release site scaffolds for rapid and precise dopamine release.

### Does Munc13 prime dopamine vesicles for fast release?

Fast and efficient neurotransmitter release relies on vesicle priming, which prepares the vesicular and plasma membranes for exocytosis and often involves vesicle attachment to the target membrane (Kaeser and Regehr, 2017). At fast synapses, RIM recruits Munc13 to active zones and activates it, and Munc13 then controls the assembly of SNARE complexes for fusion (Andrews-Zwilling et al., 2006; Betz et al., 2001; Camacho et al., 2017; Deng et al., 2011; Imig et al., 2014; Ma et al., 2013; Varoqueaux et al., 2002). For the release of modulatory transmitters, the priming mechanisms are less well understood. In some cases, they rely less on Munc13 and instead may employ alternate or additional priming pathways (Berwin et al., 1998; van de Bospoort et al., 2012; van Keimpema et al., 2017; Man et al., 2015; Renden et al., 2001). The observations that action potential triggered dopamine release requires RIM and Munc13 and is mediated by RIM zinc finger domains indicate that striatal dopamine axons employ priming mechanisms for fast release similar to conventional synapses. The finding that the dopamine system relies on these mechanisms fits well with our recent work that identified RIM as a release organizer for fast, high probability dopamine secretion (Liu et al., 2018).

Munc13 cKO^DA^ mice have altered dopamine axon and release site structure. This is unanticipated because previous studies have found normal synapse assembly in the absence of Munc13 (Augustin et al., 1999; Sigler et al., 2017; Varoqueaux et al., 2002), synaptic vesicle exocytosis (Verhage et al., 2000), or presynaptic Ca^2+^ entry (Held et al., 2020). Furthermore, loss of action potential-triggered dopamine secretion by ablation of synaptotagmin-1 (Banerjee et al., 2020) or RIM (Liu et al., 2018) does not lead to similar phenotypes. Hence, it is likely that dopamine axon structural alterations are not caused by loss of dopamine secretion itself, but Munc13 may have independent roles in dopamine axon growth and release site assembly. Effects on axon structure may be similar to a previously described role of Munc13 in the delay of growth rates of neurites in dissociated cultures and organotypic slice cultures (Broeke et al., 2010). Such roles could arise from cell autonomous functions of Munc13s, or could be mediated by knockout of Munc13-2 and Munc13-3 in surrounding cells in our experiments, for example through loss of secretion of modulatory substances important for growth. Changes in release site assembly, observed here as altered Bassoon clustering, have been described in Munc13 mutants at the fly neuromuscular junction and in cultured hippocampal neurons, where Munc13 controls the clustering of Brp or syntaxin, respectively (Böhme et al., 2016; Sakamoto et al., 2018). Hence, release site scaffolding as discovered in vivo in dopamine axons may be a Munc13 function that is shared across some secretory systems.

### Functional organization of active zone-like dopamine release sites

Initial observations suggested that RIM organizes sparse active zone-like release sites (Liu et al., 2018). However, it remained unclear if RIM operated as a scaffold at the release site or as an essential soluble or vesicle associated release factor. Here, we find that the scaffolding domains of RIM are essential, supporting the model of a scaffolded site. At classical synapses, active zone scaffolding mechanisms support three fundamental requirements for fast transmission (Biederer et al., 2017; Kaeser and Regehr, 2014; Südhof, 2012): (1) they tether Ca^2+^ channels to release sites, (2) dock vesicles to exocytotic sites, and (3) mediate the attachment and positioning of release machinery at the correct place in the target membrane. How are these functions executed to support striatal dopamine release?

Ca^2+^ channels tethering (1) for synaptic secretion is mediated by RIMs and RIM-BPs (Hibino et al., 2002; Kaeser et al., 2011; Wu et al., 2019). Several observations suggest that Ca^2+^ secretion-coupling in the dopamine system does not strongly rely on this synaptic protein complex, but may be mediated by other mechanisms. First, dopamine release is dependent on multiple Ca_V_s including Ca_V_1, Ca_V_2 and Ca_V_3 (Brimblecombe et al., 2015; Liu and Kaeser, 2019), and mechanisms that rely on direct RIM-Ca_V_2 interactions cannot explain localization of channels other than Ca_V_2s (Kaeser et al., 2011). Second, while RIM-BP may organize channels other than Ca_V_2s (Hibino et al., 2002), RIM-BP1 and -2 are dispensable for dopamine release. Third, at synapses where RIM organizes Ca_V_2s, the presence of high extracellular Ca^2+^ overrides the need for RIM (Kaeser et al., 2011, 2012), but this is not the case in the dopamine system (Liu et al., 2018). Finally, RIM-containing dopamine release sites are sparse, but Ca^2+^ entry appears to be present in all varicosities independent of the presence of RIM (Liu et al., 2018; Pereira et al., 2016). Together, these observations suggest that the RIM/RIM-BP complex is not the major or only organizer of Ca^2+^ channel complexes in dopamine axons. What other mechanisms could contribute? One possibility is that Ca_V_s are organized through transmembrane proteins rather than active zone complexes, for example neurexins which organize Ca_V_s and may drive synapse formation in cultured dopamine neurons (Ducrot et al., 2020; Luo et al., 2020). Another possibility is that α2δ proteins or β subunits drive positioning of various Ca_V_s in dopamine neurons, which could explain why subtype-specific positioning mechanism are dispensable (Held et al., 2020; Hoppa et al., 2012). While our data suggest that dopamine release does not build upon the classical Ca^2+^ secretion-coupling mechanisms, future studies should address how Ca^2+^ entry and its coupling to release ready vesicles is organized in the dopamine system.

Tethering and docking of vesicles (2) is likely important given the rapidity of dopamine release. At classical synapses, RIM and Munc13 mediate docking (Han et al., 2011; Imig et al., 2014; Kaeser et al., 2011; Wang et al., 2016; Wong et al., 2018). Technical limitations have prevented conclusive tests for vesicle docking in the dopamine system. Assessment of docking requires high pressure freezing rather than chemical fixation, which is difficult to adapt and optimize for acute brain slices from different brain regions, and dopamine-releasing varicosities are extremely sparse and difficult to identify. However, the fast kinetics and high probability of release, and requirement for Munc13 and RIM strongly suggest that dopamine vesicle docking is mediated by these proteins. Alternative or complementary attachment mechanisms could be mediated by phospholipids interactions, for example between PIP_2_ and synaptotagmin-1 (Chang et al., 2018; Jahn and Fasshauer, 2012). One interesting possibility is that both Ca^2+^ entry and vesicle-target membrane tethering, for example via synaptotagmin-1, are present in all dopamine varicosities (Banerjee et al., 2020; Pereira et al., 2016), but that priming for release only occurs in active zone containing varicosities (Liu et al., 2018; Pereira et al., 2016). This may explain why some varicosities remain silent upon stimulation despite the possible presence of RIM/Munc13 independent vesicle tethering mechanisms.

Target membrane attachment and positioning of release machinery (3) is poorly understood at synapses (Emperador-Melero and Kaeser, 2020). Proposed mechanisms include interactions with transmembrane proteins or target membrane phoshpolipds. However, strong active zone assembly phenotypes have not been reported upon disruption of any specific mechanism, for example abolishing binding to PIP_2_, or knocking out of Ca_V_2s, LAR-PTPs or neurexins (Chen et al., 2017; Held et al., 2020; de Jong et al., 2018; Sclip and Südhof, 2020). Given the dopamine secretory deficits in Liprin-α cKO^DA^ mice (Fig. 6), Liprin-α binding to LAR-PTPs (Serra-Pages et al., 1998; Serra-Pagès et al., 1995), and the dependence of dopamine release on RIM C_2_B domains which bind to Liprin-α and PIP_2_, the most parsimonious working model is that RIM C_2_B domains provide a key tethering mechanism at dopamine release sites. At synapses, secretory hotspots are strategically assembled opposed to postsynaptic receptor nanodomains, a function that may be mediated through transsynaptic cell adhesion molecules (Biederer et al., 2017; Tang et al., 2016). This model is interesting to assess in the context of the dopamine system, in which most varicosities are not directly associated with target cells through a synaptic organization (Descarries et al., 1996). One possibility is that in the dopamine system, release site localization is independent of postsynaptic receptor domains. An alternative model is that the small fraction of dopamine varicosities that is associated with postsynaptic cells (Descarries et al., 1996; Uchigashima et al., 2016) relies on such transsynaptic organization, and that only varicosities with this synaptic organization contain active zones for dopamine release, consistent with the sparsity of active zone-like assemblies and release-competent varicosities (Liu et al., 2018; Pereira et al., 2016). Future work should address the relationship between dopamine receptors and the active zone-like characterization that we describe here.

Overall, our findings establish that dopamine release sites have evolved to be fast and efficient. Scaffolding is simpler than at classical synapses based on three lines of evidence. First, functional effects of RIM deletion are stronger than at regular synapses (de Jong et al., 2018; Kaeser et al., 2011; Liu et al., 2018). Second, the scaffolds ELKS (Liu et al., 2018) and RIM-BP are entirely dispensable for dopamine release (Fig. 6). Third, the RIM C-terminal domains are essential scaffolds of dopamine release machinery (Figs. 1, 8), and Munc13 has scaffolding roles as well (Fig. 5), but at conventional synapses these structural roles are largely dispensable (Augustin et al., 1999; Deng et al., 2011; Han et al., 2011; Kaeser et al., 2011; Sigler et al., 2017; Varoqueaux et al., 2002), suggesting more redundancy. Hence, dopamine release site architecture is different from classical synapses, and relies on simple, streamlined scaffolding mechanisms.

## Acknowledgements

This work was supported by the National Institutes of Health (R01NS103484 and 5R01MH113349 to PSK, F31NS105159 to LK), the Dean’s Initiative Award for Innovation (to P.S.K.), a Harvard-MIT Joint Research Grant (to P.S.K.), a William Randolph Hearst fellowship (to A. B.), an Alice Joseph Brooks fellowship (to A. B.), a Mahoney fellowship (to A.B.), a Gordon family fellowship (to C.L.), a PhD Mobility National Grants fellowship from Xi’an Jiaotong University/China Scholarship Council (to X.C.), the European Commission (ERC Advanced Grant SynPrime, to N.B.), and Germany’s Excellence Strategy (EXC 2067/1-390729940, to N.B.). We thank J. Wang, C. Qiao, M. Han, A. Zeuch, A. Günther, and S. Beuermann for technical assistance, Dr. T.C. Südhof for the floxed RIM-BP mice, and Drs. Shan Shan Wang and Javier Emperador-Melero for help with importing the Liprin-α2 mutant mice. We acknowledge the Neurobiology Imaging Facility (supported by a NINDS P30 Core Center grant, NS072030) and the Cell Biology Microscopy Facility for availability of microscopes and advice, Dr. T. Lambert for help with 3D-SIM microscopy, Dr. M. Cicconet and the Image and Data Analysis Core (IDAC) at Harvard Medical school for help with developing TH axon analyses, the MPI-EM DNA Core Facility for mouse genotyping, the MPI-EM and Harvard Medical School animal facilities for mouse husbandry, and Dr. U. Fünfschilling and M. Schindler for blastocyst injections.

## Author contributions

Conceptualization, AB and PSK; Methodology, AB, CI, KB, LK, NL, RU, JW, XC, FB, JSR, BHC, CL, and SMW; Formal Analysis, AB, CI, KB, LK, NL, JSR, BHC, CL, SMW, NB and PSK; Investigation, AB, CI, KB, LK, NL, RU, JSR, BHC, and CL; Resources, AB, CI, KB, JW, XC, FB, SMW, NB and PSK; Writing-Original Draft, AB, CI and PSK..; Writing-Review & Editing, AB, CI, KB, LK, NL, CL, NB and PSK; Supervision, JSR, CL, SMW, NB and PSK.; Funding Acquisition NB, and PSK.

## Competing interests

The authors have no competing interests to declare.

## Methods

### Key resources table

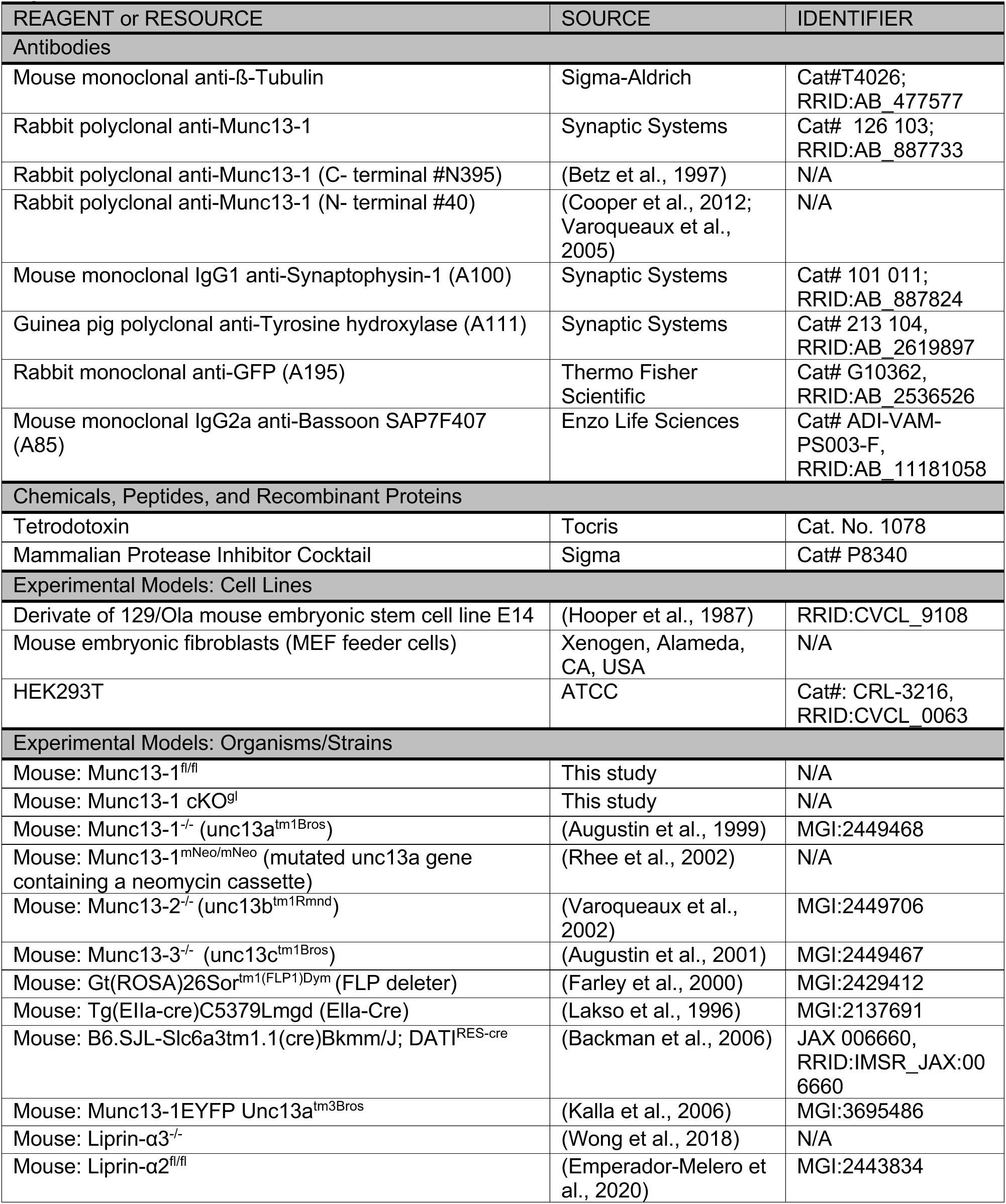
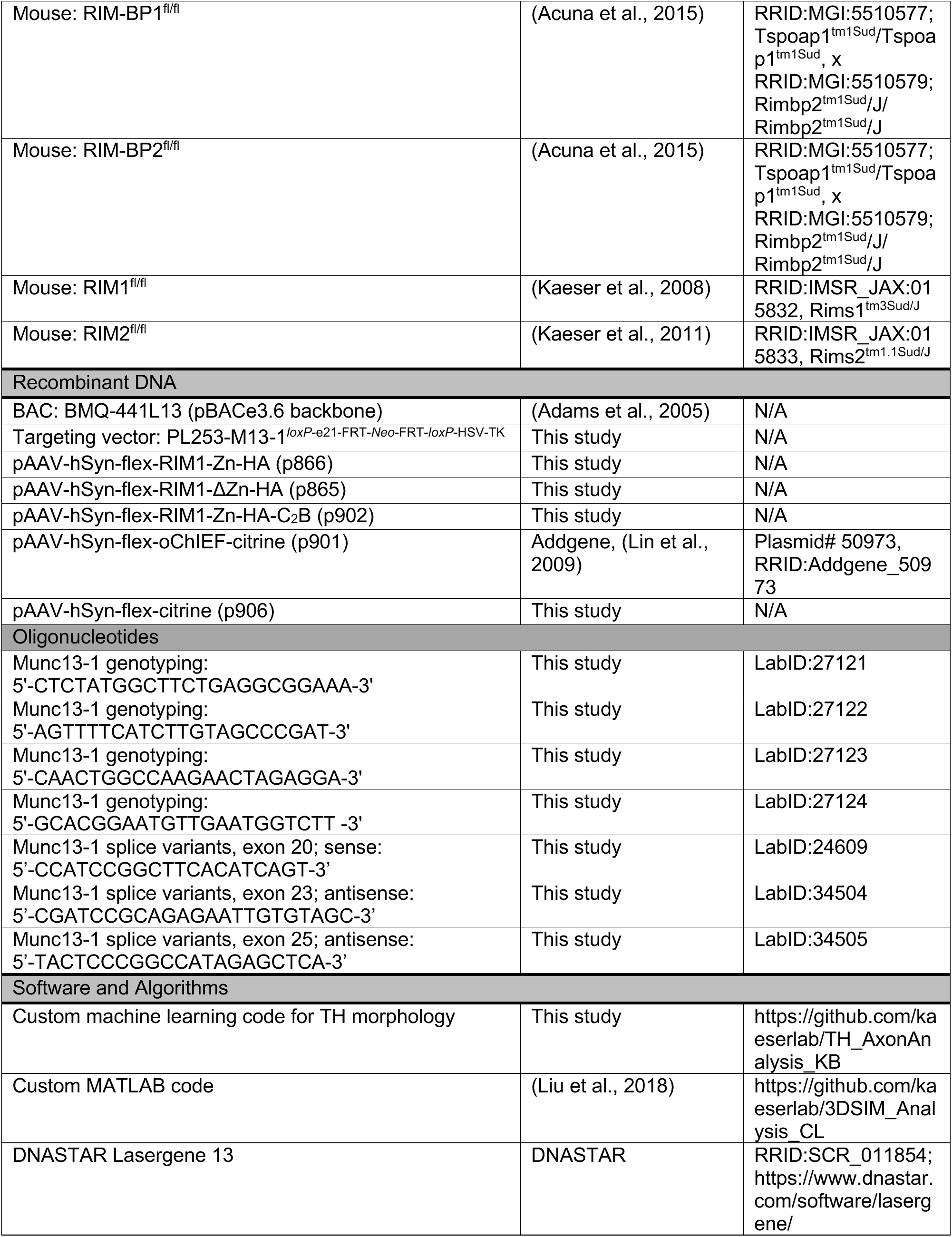
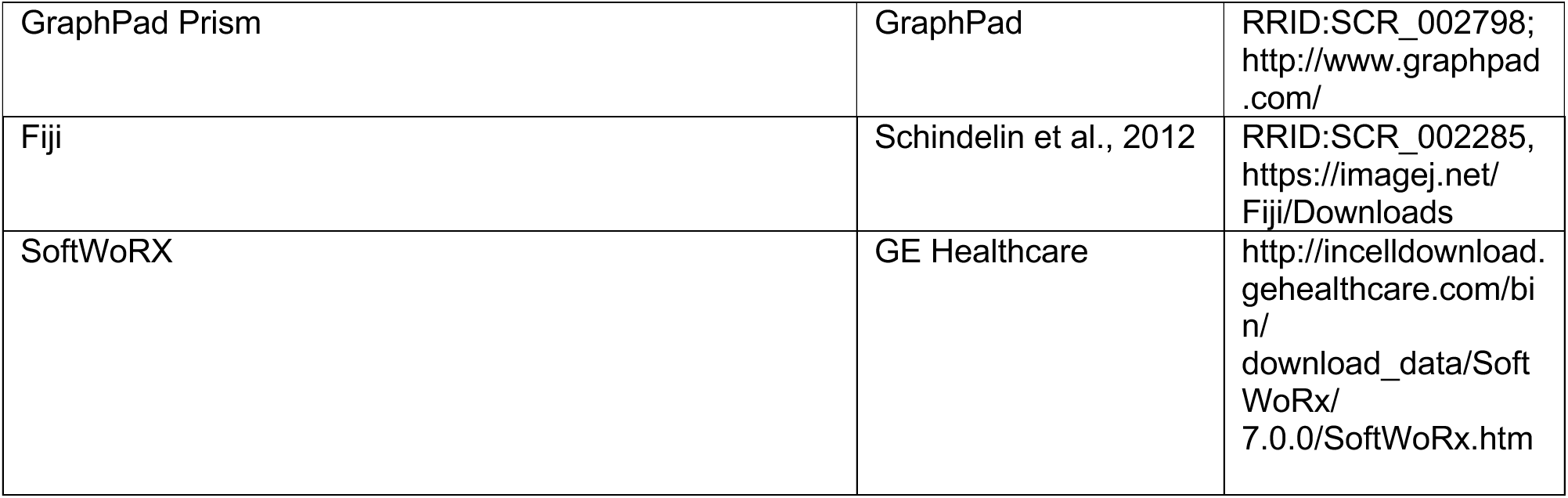

### Experimental Model and Subject Details

All animal experiments were done in accordance with approved protocols of either the Harvard University Animal Care and Use Committee, or the Niedersächsisches Landesamt für Verbraucherschutz und Lebensmittelsicherheit (LAVES; 33.19.42502-04-15/1817) and according to the European Union Directive 63/2010/EU and ETS 123. Conditional deletion of active zone proteins in dopamine neurons was performed using DAT^IRES-cre^ mice ((Backman et al., 2006); Jackson laboratories; RRID:IMSR_JAX: 006660, B6.SJL-Slc6a3^tm1.1(cre)Bkmm/^J). Unless otherwise noted, cKO^DA^ mice are mice that have two floxed alleles for active zone genes and one DAT^IRES-cre^ allele, and corresponding control mice are either siblings or age-matched mice from the same breeding colony with one floxed allele and one DAT^IRES-cre^ allele. RIM cKO^DA^ mice were generated previously (Liu et al., 2018) by breeding DAT^IRES-cre^ mice to RIM1 (RRID:IMSR_JAX:015832, Rims1^tm3Sud/J^) and RIM2 (RRID:IMSR_JAX:015833, Rims2^tm1.1Sud/J^) floxed mice. Munc13-1-EYFP mice were previously described (Kalla et al., 2006) (RRID_MGI:3695486; Unc13a^tm3Bros^), and homozygote Munc13-1-EYFP mice and unrelated age-matched control mice were used for all experiments. For removal of Munc13s in dopamine neurons (Munc13 cKO^DA^), newly generated floxed Munc13-1 mice (after crossing them to flp deleter mice, (Farley et al., 2000)) were crossed to constitutive knockout mice for Munc13-2 (Unc13b^tm1Rmnd^, RRID_MGI:2449706, (Varoqueaux et al., 2002)) and Munc13-3 (Unc13c^tm1Bros^, RRID_MGI:2449467, (Augustin et al., 2001)) and DAT^IRES-cre^ mice. Munc13 cKO^DA^ were Munc13-1^f/f^ x Munc13-2^-/-^ x Munc13-3^-/-^ x DAT^IRES-cre +/cre^. Munc13 control mice were littermates mice with Munc13-1^+/f^ x Munc13-2^+/-^ x Munc13-3^-/-^ x DAT^IRES-cre +/cre^. The Munc13-3 allele was maintained at homozygosity in breeding pairs to enable the generation of Munc13 control and cKO^DA^ siblings from the same litter. For assessment of protein content and autaptic phenotypes, Munc13-1 floxed mice were cre-recombined in the germline using EIIa-cre mice (Lakso et al., 1996), and protein content was compared to two previously established Munc13-1 knockout mouse lines (Augustin et al., 1999; Rhee et al., 2002). For deletion of RIM-BP in dopamine neurons, (RIM-BP cKO^DA^) RIM-BP1^f/f^ x RIM-BP2^f/f^ mice (RRID:MGI:5510577; Tspoap1^tm1Sud^/Tspoap1^tm1Sud^, x RRID:MGI:5510579; Rimbp2^tm1Sud^/J/ Rimbp2^tm1Sud^/J, (Acuna et al., 2015)) were obtained from Dr. T. C. Sudhof and were crossed to DAT^IRES-cre^ mice. RIM-BP cKO^DA^ were RIM-BP1^f/f^ x RIM-BP2^f/f^ x DAT^IRES-cre +/cre^ and RIM-BP control mice were littermates with RIM-BP1^+/f^ x RIM-BP2^+/f^ x DAT^IRES-cre +/cre^. For deletion of Liprin-α2 and Liprin-α3 in dopamine neurons (Liprin-α2/3 cKO^DA^), we crossed recently generated Liprin-α2^f/f^ mice (Emperador-Melero et al., 2020) to constitutive Liprin-α3 knockout mice (Wong et al., 2018) and DAT^IRES-cre^ mice. Liprin-α2/3 cKO^DA^ were Liprin-α2^f/f^ x Liprin-α3^-/-^ x DAT^IRES-cre +/cre^, and Liprin-α2/3 control mice were littermates with Liprin-α2^+/f^ x Liprin-α3^+/-^ x DAT^IRES-cre +/cre^. Given that dopamine release in control mice across experiments is similar, we conclude that heterozygosity has no strong effects on dopamine release. All mice were group housed in a 12-hr light-dark cycle with free access to water and food. All experiments with genotype comparisons were done in male and female mice, and the experimenter was blind to genotype throughout data acquisition and analysis.

### Method Details

#### Production of AAV viruses

AAVs were used to either deliver domains of active zone proteins to dopamine neurons or to stimulate dopamine neurons by inducing expression of fast channelrhodopsin oChIEF. AAV5-hSyn-FLEX-RIM1-Zn-HA, AAV5-hSyn-flex-RIM1-ΔZn-HA, and AAV5-hSyn-flex-RIM1-Zn-HA-C_2_B (which includes the linker sequence that is located between the RIM Zn finger and PDZ domains in RIM1α) were used to express specific domains of RIM1 in RIM cKO^DA^. AAV5-hSyn-flex-citrine was used as a control. For optogenetic activation of striatal dopamine fibers AAV5-hSyn-flex-oChIEF-citrine was used to drive Cre-dependent expression of oChIEF-citrine ((Lin et al., 2009), RRID:Addgene_50973). All AAVs were generated using Ca^2+^ phosphate transfection in HEK293T cells (mycoplasma free cell line from ATCC, Cat#: CRL-3216, RRID:CVCL_0063) as AAV2/5 serotypes. 72 hr after transfection, cells were collected, lysed, and viral particles were extracted and purified from the 40% layer after iodixanol gradient ultracentrifugation. Quantitative rtPCR was used to estimate the viral genomic titers (10^12^ to 10^14^ viral genome copies/ml).

#### Stereotaxic surgery

Mice (P24-P45) were anesthetized using 5% isoflurane and mounted on a stereotaxic frame. Stable anesthesia was maintained during surgery with 1.5–2% isoflurane. 1 µl of AAV viral solution was injected unilaterally into the right substantia nigra pars compacta (SNc – 0.6 mm anterior, 1.3 mm lateral of Lambda and 4.2 mm below pia) using a microinjector (PHD ULTRA syringe pump, Harvard Apparatus) at 100 nl/min. Mice were treated with post-surgical analgesia and were allowed to recover for at least 21 days prior to electrophysiology. Stereotaxic procedures were performed according to protocols approved by the Harvard University Animal Care and Use Committee.

#### Electrophysiology in brain slices

Recordings in acute brain slices were performed in the dorsolateral striatum as described before (Banerjee et al., 2020; Liu et al., 2018). Male and female mice at 42-191 days of age were anesthetized with isoflurane and decapitated. 250 µm thick sagittal brain sections containing dorsal striatum were cut on a vibratome (Leica, VT1200s) using ice-cold sucrose-based cutting solution with (in mM): 75 sucrose, 75 NaCl, 26.2 NaHCO_3_, 1 NaH_2_PO_4_, 1 sodium ascorbate, 2.5 KCl, 7.5 MgSO_4_, 12 glucose, 1 myo-inositol, 3 sodium pyruvate, pH 7.4, 300–310 mOsm. Slices were incubated in incubation solution bubbled with 95% O_2_ and 5% CO_2_ containing (in mM): 126 NaCl, 26.2 NaHCO_3_, 1 NaH_2_PO_4_, 2.5 KCl, 1 sodium ascorbate, 3 sodium pyruvate, 1.3 MgSO_4_, 2 CaCl_2_, 12 glucose, 1 myo-inositol (pH 7.4, 305–310 mOsm) at room temperature for 1 hour. All recordings were done at 34–36°C, and slices were continuously perfused with artificial cerebrospinal fluid (ACSF) at 3–4 ml/min bubbled with 95% O_2_ and 5% CO_2_. ACSF contained (in mM): 126 NaCl, 26.2 NaHCO_3_, 2.5 KCl, 2 CaCl_2_ (unless noted otherwise), 1.3 MgSO_4_, 1 NaH_2_PO_4_, 12 glucose, pH 7.4, 300–310 mOsm. Recordings were completed within 5 hr of slicing. The experimenter was blind to genotype throughout recording and data analyses. All data acquisition and analyses for electrophysiology were done using pClamp10 (Clampex, Axon Instruments).

For <u>carbon fiber amperometry</u>, carbon fiber microelectrodes (CFEs, 7 µm diameter, 200–350 µm long) were custom-made by inserting carbon fiber filaments (Goodfellow) into glass capillaries. On the day of recording, each new CFE was calibrated before use by puffing freshly made dopamine solutions of increasing concentrations (0, 1, 5, 10, 20 µM) in ACSF for 10 s. The currents for each concentration of dopamine were plotted against the dopamine concentration and only CFEs with a linear relationship were used. For all genotype comparisons, each control and cKO^DA^ littermate pair was recorded in an interleafed manner on the same day using the same carbon fiber. For RIM cKO^DA^ rescue experiments, an uninjected control mouse was first recorded to establish stable carbon fiber recordings followed by inter-leafed recordings from two injected littermate RIM cKO^DA^ mice. CFEs were slowly inserted 20–60 µm below the slice at dorsolateral striatum and were held at 600 mV to record dopamine release. Signals were sampled at 10 kHz and low-pass filtered at 400 Hz. Dopamine release in dorsolateral striatum was evoked by electrical or optogenetic stimulation every 2 min.

<u>Electrical stimulation</u> was applied through ACSF filled glass pipette (tip diameter 3–5 µm) connected to a linear stimulus isolator (A395, World Precision Instruments) was used to deliver monopolar electrical stimulation (10–90 µA) to striatal slices and elicit dopamine release. The stimulation pipette was kept at 20–30 µm below the slice surface in dorsolateral striatum and 100–120 µm away from the tip of CFE. A biphasic wave (0.25 µs in each phase) was applied to evoke dopamine release. Electrical stimulation was delivered either as a single stimulus or 10 Hz trains of 90 µA or single stimuli of increasing intensities 10-90 µA. <u>Optogenetic stimulation</u> was used to evoke dopamine release in areas in the dorsolateral striatum with uniform citrine fluorescence. Optogenetic stimulation was applied as ten pulses of 470 nm light (each of 1 ms duration) as a 10 Hz train at the recording site through a 60 x objective by a light-emitting diode (Cool LED pE4000). Optogenetic stimulation was applied every 2 min for all dopamine release measurements or every 10 seconds for extracellular field recordings. For extracellular field recordings, optogenetic stimulation was either applied as ten pulses of 470 nm light at 10 Hz or as forty pulses at 40 Hz. 1 µM TTX (Tocris, Catalogue No.# 1078) was used to inhibit sodium channels and action potential firing-dependent dopamine release.

<u>KCl stimulation</u> was done using a solution containing (in mM) 100 KCl, 1.3 MgSO_4_, 50 NaCl, 2 CaCl_2_ (unless mentioned otherwise), 12 glucose, 10 HEPES (pH 7.3, 300–310 mOsm). KCl was puffed onto dorsolateral striatum for 10 seconds at 9 µl/s using a syringe pump (World Precision Instruments) and recordings were in ACSF (unless mentioned otherwise). The peak amplitude of the dopamine response was quantified, and the area under the curve was measured from start of KCl puff for 50 s. Only one KCl puff was applied per slice except for Figs. S1G, S1H. For assessing the Ca^2+^-dependence of KCl evoked dopamine release (Figs. S1G, S1H), the 0 mM Ca^2+^ KCl puff solution contained (in mM): 100 KCl, 3.3 MgSO_4_, 50 NaCl, 1 mM EGTA, 12 glucose, 10 HEPES (pH 7.3, 300–310 mOsm), and was puffed onto the recording site. Slices were perfused with 0 mM Ca^2+^ ACSF containing (in mM): 126 NaCl, 3.3 MgSO_4_, 26.2 NaHCO_3_, 2.5 KCl, 1 EGTA, 1 NaH_2_PO_4_, 12 glucose, (pH 7.4, 300–310 mOsm) during recording. Slices were either incubated in 2 mM Ca^2+^ (regular) ACSF or 0 mM Ca^2+^ ACSF for at least 1 h prior to start of recording. Slices incubated in 2 mM Ca^2+^ containing ACSF received two 2 KCl puffs separated by an interval of 15 mins, and both KCl puff solutions contained 2 mM Ca^2+^ and no EGTA. Slices incubated in 0 mM Ca^2+^ ACSF, the first KCl puff was done with 0 mM Ca^2+^ KCl puff and recording was done in 0 mM Ca^2+^ ACSF. Slices were then perfused for 15 min with 2 mM Ca^2+^ ACSF and a second KCl puff with 2 mM Ca^2+^ followed.

<u>Extracellular field potential recordings</u> were used to record dopamine axon action potential firing as described before (Banerjee et al., 2020; Liu et al., 2018) and were performed with ACSF filled glass pipettes (2–3 µm tip diameter) placed 20–60 µm below the slice surface in areas of dorsolateral striatum with uniform citrine fluorescence. Optogenetic stimulation was applied as a 10 Hz train or a 40 Hz train every 10 s and 100 sweeps were averaged for quantification. Sodium channels were blocked using 1 µM TTX (Tocris, Catalogue No.# 1078) and extracellular potentials evoked by 10 Hz trains were recorded before and after TTX. To quantify the reduction by TTX, the amplitude evoked by the first stimulus of the 10 Hz train before and after TTX was analyzed.

#### Immunostaining of brain sections

Male and female mice (104–205 days old) were deeply anesthetized with 5% isoflurane. Transcardial perfusion was performed with ice-cold 30–50 ml phosphate buffer saline (PBS) and brains were fixed by perfusion with 50 ml of 4% paraformaldehyde in PBS (4% PFA) at 4°C. Brains were dissected out and incubated in 4% PFA for 12–16 hr followed by dehydration in 30% sucrose + 0.1% sodium azide in PBS overnight or until they sank to the bottom of the tube. 20 µm thick coronal striatal sections were cut using a vibratome (Leica, VT1000s) in ice-cold PBS. Next, antigen retrieval was performed by incubating slices overnight at 60°C in 150 mM NaCl, 0.05% Tween 20, 1 mM EDTA, 10 mM Tris Base, pH 9.0. Slices were washed in PBS for 10 min and incubated in PBS containing 10% goat serum and 0.25% Triton X-100 (PBST) for 1 hr at room temperature. Slices were incubated with primary antibodies for 8-12 hr at 4°C, and the following primary antibodies were used in PBST: mouse monoclonal IgG2a anti-Bassoon (1:500, A85, RRID:AB_11181058), guinea pig polyclonal anti-TH (1:1000, A111, RRID:AB_2619897) and rabbit monoclonal anti-GFP (1:2000, A195, RRID:AB_2536526). Slices were washed thrice in PBST each for 10 mins and incubated in secondary antibodies for 2 hr at room temperature in PBST. Secondary antibodies used were goat anti-mouse IgG2a Alexa 488 (1:500, S8, RRID:AB_2535771), goat-anti guinea pig Alexa 568 (1:500, S27, RRID:AB_2534119) and goat anti-rabbit Alexa 488 (1:500, S5, RRID:AB_2576217). Sections were washed thrice in PBST each for 10 mins to wash off excess secondary antibodies and mounted on Poly-D-lysine coated #1.5 cover glasses (GG-18–1.5-pdl, neuVitro) with H-1000 mounting medium (Vectashield). At all times during perfusion, staining and mounting, the experimenter was blind to the genotype of the mice.

#### 3D-SIM image acquisition and analysis

Image acquisition and analyses were done essentially as described before (Banerjee et al., 2020; Liu et al., 2018) using a DeltaVision OMX V4 Blaze structured illumination microscope (GE Healthcare) with a 60 x 1.42 N.A. oil immersion objective and Edge 5.5 sCMOS cameras (PCO) for each channel. Z stacks were acquired in the dorsolateral striatum with a 125 nm step size and 15 raw images were obtained per plane (five phases, three angles). Immersion oil matching was used to minimize spherical aberration. A control slide with TH axonal staining in red and green fluorophores was used to measure lateral shift between green and red channels. A calibration image was generated from this control slide and all images were reconstructed using this calibration to reduce lateral shifts between fluorophores. All 3D-SIM raw images were aligned and reconstructed to obtain superresolved images using the image registration function in softWoRx. Image volumes (40 x 40 x 6 µm^3^) were acquired from 7 to 8 regions within the dorsolateral striatum in 4-5 coronal sections for each animal. For detection of Munc13-1 within TH axons, anti-GFP antibodies were used to visualize Munc13-1 in mice in homozygote mice in which Munc13-1 is endogenously tagged with EYFP mice (Kalla et al., 2006), and age-matched wild type mice were used as negative controls. The intensity range for Munc13-1 puncta was determined from reconstructed images, and multiple intensity thresholds within this range were used to generate masks of Munc13-1 puncta, and settings in which Munc13-1 masks best matched original images irrespective of their relationship to TH were chosen for quantification by an investigator blind to the genotype of the mice, and the same thresholds were then used for the full dataset. For image analyses, regions of interest (ROIs) ranging from 20 x 20 x 2.5 µm^3^ to 25 x 25 x 2.5 µm^3^ were selected manually in each z-stack of Munc13-1-EYFP and wild type images. To characterize TH and Munc13-1 signals, size thresholds (0.04–20 µm^3^ for TH axons, 0.003–0.04 µm^3^ for Munc13-1) were applied. For detection of bassoon within TH axons in Munc13 control and Munc13 cKO^DA^ mice, ROIs were generated using Otsu intensity and size thresholding parameters (0.04–20 µm^3^ for TH axons, 0.003–0.04 µm^3^ for Bassoon). The overlap of Munc13-1 and bassoon with TH axon was quantified using a custom written MATLAB code (Liu et al., 2018) (available at https://github.com/kaeserlab/3DSIM_Analysis_CL or https://github.com/hmslcl/3D_SIM_analysis_HMS_Kaeser-lab_CL). The volume occupied by TH for each image was quantified and divided by the total image volume. This was followed by skeletionization of TH signals using 3D Gaussian filtering and a homotypic thinning algorithm to calculate TH axon length. Munc13-1 and Bassoon objects were considered to be within TH when there was > 40% overlap, as established before (Liu et al., 2018). For quantification of Munc13-1 clusters, density and volume occupied by Munc13-1 objects per image volume were calculated. Density and volume of Munc13-1 objects within TH was quantified, and compared to the average of the same parameters after shuffling each object within 1 x 1 x 1 µm^3^ for 1000 times. For quantification of bassoon clusters in Munc13 control and cKO^DA^, the density and volume of bassoon clusters showing >40% overlap with TH axon was quantified. For quantitative assessment of TH axon shape, a custom code was generated (available at https://github.com/kaeserlab/TH_AxonAnalysis_KB). Briefly, a machine learning model was trained with annotation of 15 images per genotype for detection of TH positive objects vs background. All images were next processed in a 3D-smoothing operation followed by thresholding on blurred probability maps. The central axis, the radius and total surface area for each axonal segment was computed. The proportion of surface area at a specific distance from the central axis, in 0.04 µm increments, were compared between Munc13 control and cKO^DA^. Sample images were generated using Imaris 9.0.2 (Oxford Instruments) from masked images obtained from the custom analysis code. Adjustments of contrast, intensity and surface rendering were done identically for each condition for illustration, but after quantification. For all 3D-SIM data acquisition and analyses, the experimenter was blind to the genotype of the mice.

#### Generation of conditional Munc13-1 knockout mice

Mice with a floxed exon 21 of the Munc13-1 gene were generated by homologous recombination in 129/Ola embryonic stem (ES) cells according to standard protocols. The targeting vector was generated by recombineering and subcloning from a bacterial artificial chromosome (BAC; BMQ-441L13) of a 129SV ES cell DNA BAC library, and a Herpes Simplex Virus Thymidine kinase (HSV-TK) and neomycin resistance cassette were used as positive and negative selection markers, respectively. ES cell clones were analyzed by Southern blotting of HindIII-digested genomic DNA, and positive ES cell clones were injected into blastocysts to obtain chimeric mice. FLP deleter mice (Farley et al., 2000) were used to generate the Munc13-1 floxed mice. Genotyping was performed by PCR using the following reactions: Munc13-1 wild type allele with 27123 + 27122 yielding a 149 bp band; Munc13-1 knock-in allele (ki) after homologous recombination with 27121 + 27122 yielding a 196 bp band; Munc13-1 floxed allele after FLP-mediated excision of neomycin resistance cassette with 27123 + 27122 yielding a 253 bp band; Munc13-1 cKO allele after Cre mediated recombination with 27124 + 27122 yielding a 209 bp band. For whole brain Western blotting and electrophysiological assessment of synaptic transmission in autaptic neurons, Munc13-1 floxed mice were crossed to Ella-Cre mice for germline recombination (Lakso et al., 1996) to produce constitutive Munc13-1 cKO^gl^ mice. For rtPCR, total RNA was isolated from wild type and Munc13-1 cKO^gl^ P0 mouse brains using the Direct-zol RNA Miniprep Kit and reverse transcribed into double-stranded-cDNA by the SuperScript™ Double-Stranded cDNA Synthesis Kit. PCR reactions were performed using the proofreading VELOCITY DNA polymerase and the primer pairs 24609/34504 (exon 20-23: wt: 450 bp; Δe21: 286 bp; Δe21+22: 135 bp) and 24609/34505 (exon 20-25: wt: 679 bp; Δe21: 515 bp; Δe21+22: 364 bp). The sequences of the PCR products obtained from the different splice variants were verified by directly sequencing the purified PCR products and after cloning them into a TOPO TA cloning vector.

#### Immunoblotting

Western blots were used to estimate Munc13-1 levels in whole brain homogenates from P0 Munc13-1 wild type Munc13-1^f/f^ littermates, and 5 µg from each sample were loaded onto a 3-8% Tris-acetate gradient gel. To compare Munc13 expression in whole brain homogenates of different Munc13 mouse mutants, 20 µg protein per sample was separated on a 7.5 % gel. Residual Munc13-1 expression was estimated in whole brain homogenates from Munc13-1 cKO^gl^ animals and Munc13-1 wild-type littermates. 20 µg from Munc13-1 cKO^gl^ and varying concentrations (0.7 µg, 1.3 µg, 2 µg) from Munc13-1 wild type homogenates were loaded onto each lane of a 3-8% Tris-acetate gel. After the transfer, protein bound to nitrocellulose membranes was visualized with MemCode. Membranes were destained and then washed with TBS buffer (10 mM Tris-HCl, 150 mM NaCl, pH 7.5), incubated with blocking buffer (TBS, 5% (w/v) milk powder, 5% (v/v) goat serum, 0.1% (v/v) Tween-20) for 30 min, incubated with primary antibodies for 1 h at room temperature, washed in TBS-T buffer (TBS, 0.1% (v/v) Tween-20), and incubated with secondary antibodies conjugated to horseradish peroxidase (goat anti-Mouse IgG (H+L), Jackson ImmunoResearch, Cat#115-035-146, RRID:AB_2307392; goat anti-Rabbit IgG (H+L), Jackson ImmunoResearch, Cat#111-035-144, RRID: AB_2307391). After several washing steps in TBS-T and TBS, immunoreactive bands were visualized with an enhanced chemiluminescence (ECL) detection system on film. The following primary antibodies were used: anti-Munc13-1 [#40, rabbit polyclonal, N-terminal, 1:1,000 (Cooper et al., 2012; Varoqueaux et al., 2005); #N395, rabbit polyclonal, C-terminal, 1:250 (Betz et al., 1997); SySy #126 103, rabbit polyclonal, N-terminal 1:1,000, RRID:AB_887733], anti-ß-Tubulin (Sigma-Aldrich, #T4026, mouse monoclonal, 1:5,000, RRID:AB_477577). For estimation of protein amounts, films and stained nitrocellulose membranes were scanned and analyzed using ImageJ (Schindelin et al., 2012). Protein levels were normalized to the total protein amount in the sample as measured by the MemCode stain using the tracing tool. Protein bands from films were manually outlined and the signal intensity was measured. The signal intensity of the band was normalized to the total protein in the respective lane. For experiments estimating residual Munc13-1 expression, the normalized averages of the 0.7 µg/lane and 1.3 µg/lane Munc13-1 wild type samples were compared to 20 µg/lane Munc13-1 cKO^gl^ samples.

#### Generation of autaptic mouse hippocampal neuron cultures

Astroglial feeder monolayer cell cultures were generated from wild-type (C57/N) postnatal day (P0) mouse cortices according to a previously published protocol (Burgalossi et al., 2012). Primary neuron cultures were prepared from neonatal mouse brains that were dissected at P0 in ice-cold Hank’s Balanced Salt Solution (HBSS). Both hippocampi were removed and transferred into 500 µl of prewarmed Papain solution (DMEM supplemented with 20 units/ml papain; 0.2 mg/ml cysteine; 1 mM CaCl_2_; 0.5 mM EDTA) and incubated for 60 min at 37°C. The digestion of the hippocampi was terminated by incubating the tissue for 15 min in inactivation solution (DMEM supplemented with 2.5 mg/ml BSA; 2.5 mg/ml trypsin inhibitor; and 10% (v/v) FBS). After two medium washes, neurons were dissociated and seeded onto microdot astrocyte feeder islands on glass coverslips (4,000 cells/6 well for electrophysiology, S2E-M). Neurons were maintained in culture medium (Neurobasal-A medium supplemented with 1x B27, 2 mM Glutamax, and 100 units/ml penicillin/streptomycin) at 37°C and 5% CO_2_.

#### Electrophysiology on neurons in autaptic hippocampal cultures

Autaptic hippocampal neurons from Munc13-1 wild type and Munc13-1 cKO^gl^ littermate mice were whole-cell voltage clamped at DIV 13-16. Neurons were recorded at room temperature in an external bath solution containing (in mM): 140 mM NaCl, 2.4 mM KCl, 10 mM HEPES, 10 mM glucose, 4 mM CaCl_2_, and 4 mM MgCl_2_ (320 mOsm/l) pH 7.4. Patch pipettes (2.5-3.8 MΩ) were filled with internal solution containing (in mM): 136 mM KCl, 17.8 mM HEPES, 1 mM EGTA, 4.6 mM MgCl_2_, 4 mM NaATP, 0.3 mM Na_2_GTP, 15 mM creatine phosphate, and 5 U/ml phosphocreatine kinase, pH 7.4, 315–320 mOsm. Excitatory postsynaptic currents (EPSCs) were evoked in patched neurons by a 2 ms depolarization to 0 mV. The peak amplitudes for all responses recorded in 10 Hz train across both genotypes was normalized to the mean initial EPSC amplitude of Munc13-1 control. Miniature EPSCs (mEPSCs) were recorded in presence of 300 nM TTX to inhibit action potential firing. 500 mM sucrose was puffed for 7 seconds to estimate the readily-releasable pool of synaptic vesicles.

#### Microdialysis

Microdialysis was performed as described before (Banerjee et al., 2020; Liu et al., 2018). The microdialysis probes (6 kDa MW cut-off, CMA 11, Harvard Apparatus, Catalogue# CMA8309581) were calibrated with freshly made dopamine solutions (0, 100 and 200 nM) dissolved in ACSF before each experiment. After probe calibration, male and female Munc13 control and cKO^DA^ mice (55–96 days old) were anesthetized using 1.5% isoflurane and the probe was inserted into dorsal striatum (coordinates: 1.0 mm anterior, 2.0 mm lateral of bregma, and 3.3 mm below pia) using stereotaxy. A fresh probe was used for each Munc13 control and cKO^DA^ mouse. The microdialysis probe was continuously perfused with ACSF containing (in mM): 155 NaCl, 1.2 CaCl_2_ 1.2 MgCl_2_, 2.5 KCl and 5 glucose at a speed of 1 µl/min. Dialysates from dorsal striatum were collected every 15 min and the concentration of extracellular dopamine was measured using HPLC (HTEC-510, Amuza Inc) connected to an electrochemical detector (Eicom). Data during the first 75 min were not plotted because during this time window dopamine levels stabilize after surgery. Average dopamine levels from the 76th - 120th min of Munc13 control mice were used to normalize all dopamine values for both genotypes. 10 µM TTX dissolved in ACSF was applied using reverse dialysis starting at 121 min to inhibit firing of dopamine axons as described before (Banerjee et al., 2020; Liu et al., 2018). For all microdialysis data acquisition and analyses, the experimenter was blind to the genotype of the mice, and experiments were performed according to approved protocols of the Harvard University Animal Care and Use Committee.

#### Striatal synaptosome preparation and immunostaining

Striatal synaptosome preparations were performed as previously described (Banerjee et al., 2020; Liu et al., 2018). Munc13 control and cKO^DA^ mice (P42-73) were deeply anesthetized using isoflurane, decapitated, and the brains were harvested into ice-cold PBS. Dorsal striata were dissected out and placed into a pre-cooled, detergent-free glass-Teflon homogenizer filled with 1 ml of ice-cold homogenizing buffer containing (in mM): 4 4-(2-hydroxyethyl)-1-piperazineethanesulfonic acid (HEPES), 320 sucrose, pH 7.4, and 1x of a mammalian protease inhibitor cocktail. The tissue was homogenized with 12 strokes with a detergent-free ice-cold glass-teflon homogenizer. Next, 1 ml of homogenizing buffer was added to the striatal homogenate and it was centrifuged at 1,000 g for 10 min at 4°C. The supernatant (S1) was collected and centrifuged at 12,500 g for 15 min at 4°C. The supernatant (S2) was removed and the pellet (P2) was re-homogenized in 1 ml homogenizing buffer with 6 strokes. A sucrose density gradient was prepared with 5 ml of both 0.8 M and 1.2 M sucrose in thin wall ultracentrifugation tubes (Beckman Coulter, Cat # 344059). P2 homogenate was mixed with 1 ml of homogenizing buffer, and 1.5 ml of this was added to the top of the sucrose gradient and was centrifuged at 69,150 x g for 70 min at 4°C (SW 41 Ti Swinging-Bucket Rotor, Beckman Coulter, Cat. # 331362). 1–1.5 ml of the synaptosome layer was collected from the interface of the two sucrose layers. Synaptosomes were diluted 20–30 times in homogenizing buffer and spun (4000 x g, 10 min) onto Poly-D-lysine coated #1.5 coverslips at 4°C. Excess homogenizing buffer was pipetted out and synaptosomes were fixed using 4% PFA in PBS for 20 min at 4°C. Coverslips were incubated in 3% bovine serum albumin + 0.1% Triton X-100 in PBS at room temperature for 45 min to block non-specific binding and allow for permeabilization. Primary antibody staining was done for 12 hr at 4°C, followed by three washes for 15 min each. The primary antibodies used were: mouse monoclonal IgG2a anti-Bassoon (1:1000, A85, RRID:AB_11181058), mouse monoclonal IgG1 anti-Synaptophysin-1 (1:500, A100, RRID:AB_887824) and guinea pig polyclonal anti-TH (1:1000, A111, RRID:AB_2619897). Secondary antibody staining was done for 2 hr at room temperature in blocking solution followed by three washes each for 15 min. The secondary antibodies were: goat anti-mouse IgG2a Alexa 488 (1:500, S8, RRID:AB_2535771), goat anti-rabbit Alexa 555 (1:500, S22, RRID:AB_2535849), and goat anti-guinea pig Alexa 633 (1:500, S34, RRID:AB_2535757).

#### Confocal microscopy and image analysis of striatal synaptosomes

Single optical sections of striatal synaptosomes plated on coverslips (105 x 105 µm^2^) stained for Bassoon (detected via Alexa 488), Synaptophysin-1 (detected via Alexa-555) and TH (detected via Alexa 633) were imaged with an oil immersion 60 x objective and 1.5 x optical zoom using an Olympus FV1000 confocal microscope. For quantification, raw confocal images were analyzed in a custom MATLAB program (Liu et al., 2018) (available at https://github.com/hmslcl/3D_SIM_analysis_HMS_Kaeser-lab_CL). A total of 300-700 synaptosomes were detected per image using Otsu intensity thresholds and size thresholds (0.2–1 µm^2^ for TH and 0.15-2 µm^2^ for synaptic markers). These threshold settings were identical for each image across Munc13 control and cKO^DA^ for detection of Synaptophysin-positive (Syp^+^) and TH-positive (TH^+^) ROIs in each image, which was used to generate single and double positive ROIs (Syp^+^TH^+^). For Synaptophysin-positive TH-negative ROIs (Syp^+^TH^-^), Synaptophysin^+^ ROIs which had a TH signal less than the average intensity of all pixels in the image were designated as TH-negative (TH^-^) and were marked as (Syp^+^TH^-^) ROIs. Bassoon intensities within Syp^+^TH^-^, Syp^+^TH^+^ ROIs (Fig. 5) or within all TH^+^ ROIs (Fig. S5) were quantified and frequency distribution histograms were plotted, and Synaptophysin intensity within all TH^+^ ROIs was quantified. Sample images were generated in Fiji with identical adjustments of brightness and contrast for Munc13 control and cKO^DA^, but image quantification was performed before these adjustments. Image acquisition and quantification was performed by an experimenter blind to the genotype.

#### Statistics

All data were expressed in mean ± SEM (except for 5J, mean ± SD) and all statistics were performed in Graphpad Prism 9. Unpaired t-tests were used in Figs. 4D inset, 4E inset, 5B, 5C, 5G, 5H, 5K, 5L, 6C inset, 7C inset, S5B; Mann Whitney tests were used in Figs. 1D, 1F, 1I, 1K, 1M, 1O, 1Q, 1S, 2C, 2D, 2E, 2F, 2G, 3G, 3H, 6E, 6F, 7E, 7F, 8C, 8E, S3B, S3C, S3E, S3G; Wilcoxon tests were used in Figs. 2H, 2I; one-way ANOVA followed by Sidak’s multiple comparisons test were used in Figs. 3D, 5D, S1H, S4D; one-way ANOVA followed by Dunnett’s multiple comparisons test in Figs. S8C, S8D; two-way ANOVA followed by Sidak’s multiple comparisons test were used in Figs. 3E, 4B, 4D, 4E, 6C, 7C, S3I, S4E, S4G, S6B, S6D, S7B, S7D; Kolmogorov-Smirnov test were used in Figs. 5E, S5C and Chi-square test was used in Fig. 5J.

---

**Figure S1.**
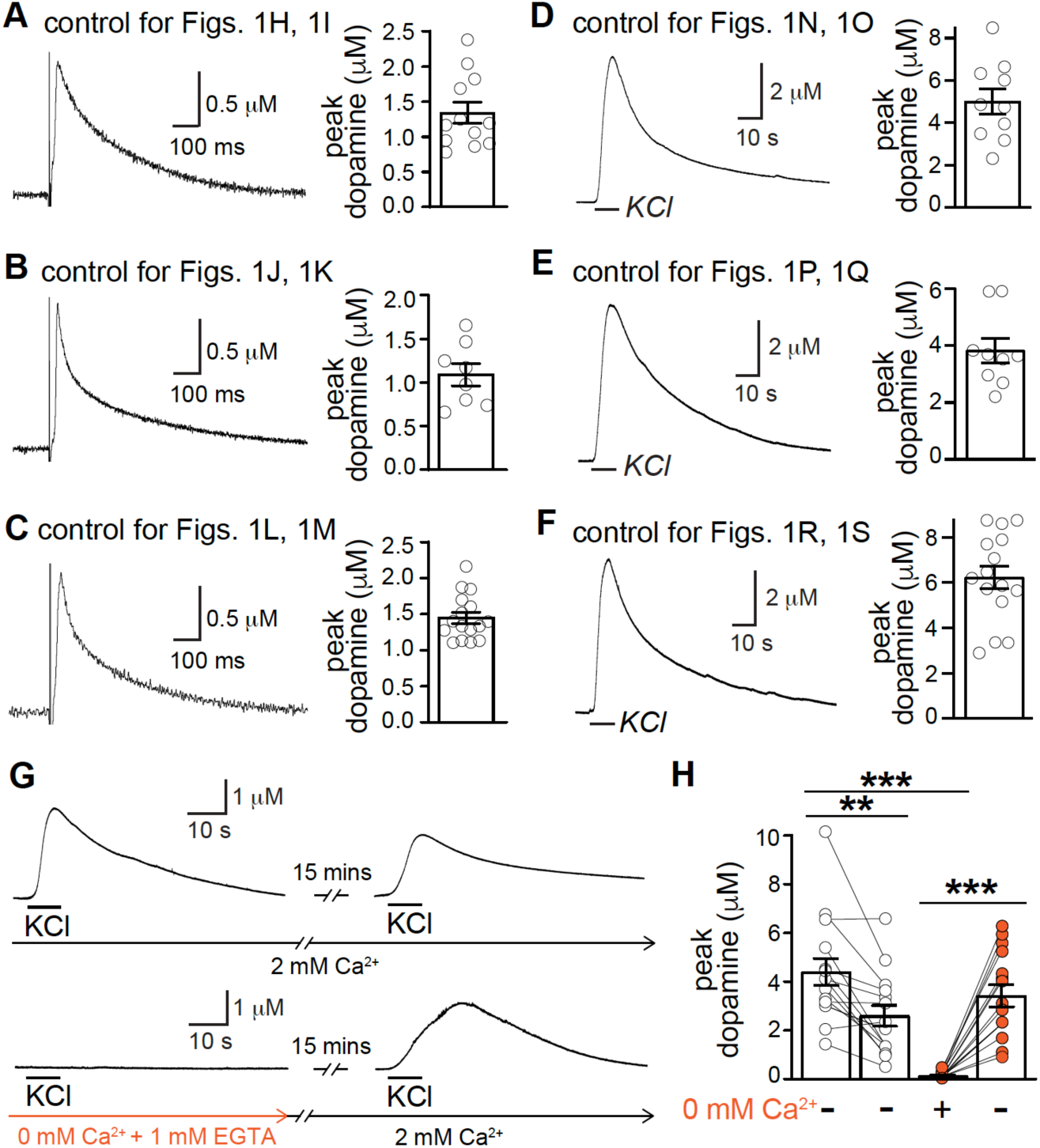
Additional data for RIM rescue experiments and Ca^2+^-dependence of KCl-triggered release, related to Fig. 1 **(A-C)** Sample traces (single sweeps) and quantification of dopamine release evoked by a 90 µA electrical stimulus in unrelated control slices of dorsolateral striatum that were used for each experiment shown in Fig. 1 to establish stable amperometric recordings during rescue experiments. n = 12 slices/4 mice in A; 8/3 in B; 16/4 in C. **(D-F)** Same as A-C but for a local 100 mM puff of KCl. n = 10/4 in D; 9/3 in E; 15/4 in F. **(G-H)** Sample traces (G) and quantification of peak dopamine amplitudes (H) evoked by 2 consecutive 100 mM KCl puffs at an interval of 15 min. Slices were incubated either in 2 mM Ca^2+^ throughout the experiment or in 0 mM Ca^2+^ + 1 mM EGTA containing ACSF for the first pulse and then in 2 mM Ca^2+^ for the second pulse. 2 mM Ca^2+^ = 15 slices/4 mice; 0 mM Ca^2+^ + 1 mM EGTA = 15/4. Data are mean ± SEM, ** p < 0.01, *** p < 0.001 as assessed by one-way ANOVA followed Sidak’s multiple comparisons test in H.

**Figure S2.**
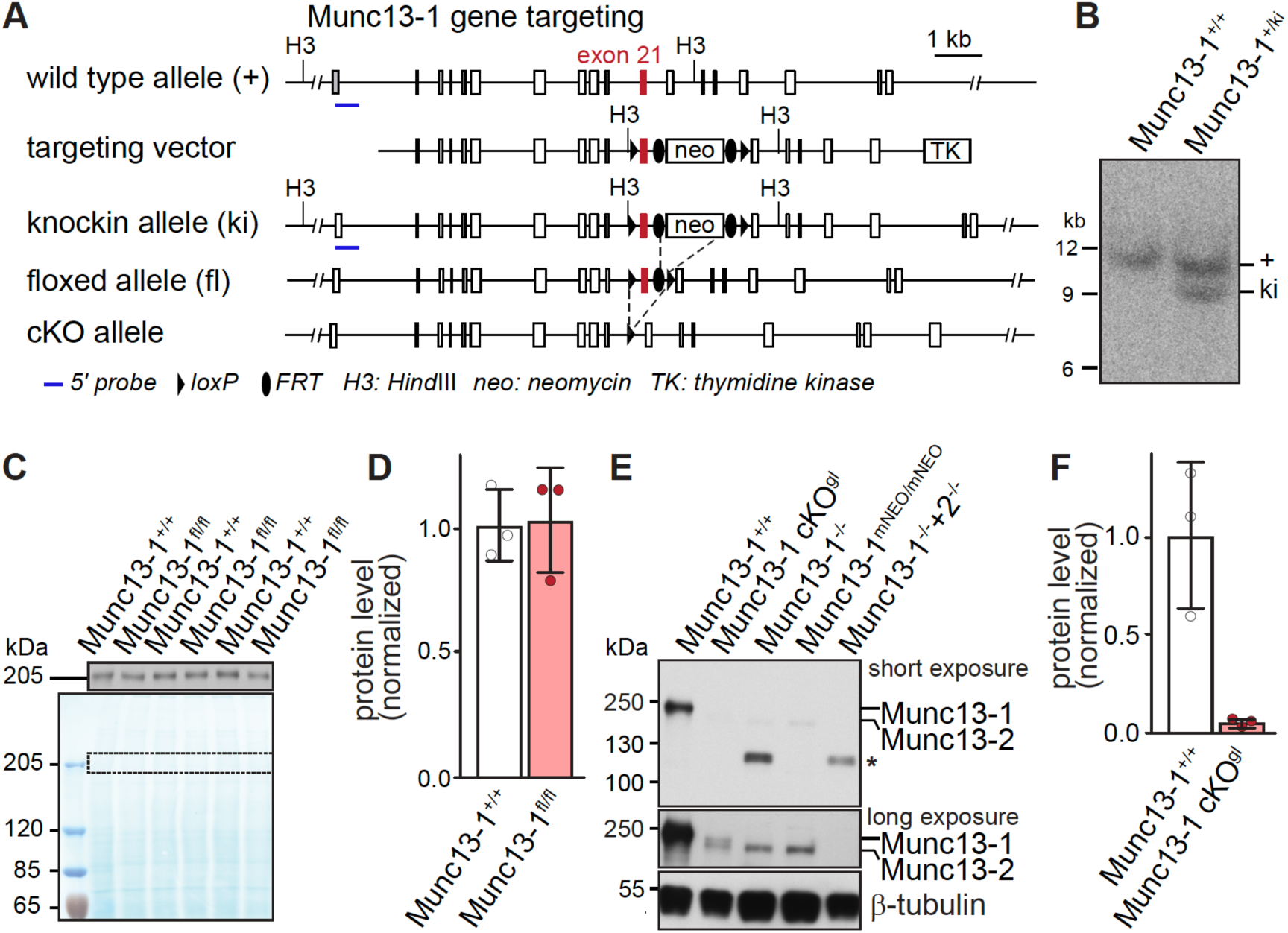
Generation of Munc13-1 floxed mice, related to Fig. 3 **(A)** Schematic of Munc13-1 wild type allele, the targeting vector, the knock-in allele (ki) after homologous recombination, the floxed allele after FLP-mediated excision of the neomycin resistance cassette (neo), and the Munc13-1 cKO allele after Cre-mediated recombination. The targeted exon 21 is indicated in red. A 5’ probe (blue bars) was used for Southern blot analysis in B. **(B)** 5’ probe Southern blot hybridization after *Hind*III-digestion of genomic DNA purified from a positive embryonic stem (ES) cell clone produced 9.8 kb Munc13-1 wild type and 8.4 kb Munc13-1 ki band. **(C, D)** Western blot (C) and estimation of Munc13-1 levels (D) in P0 whole brain homogenates from wild type (Munc13-1^+/+^) and homozygote floxed (Munc13-1^fl/fl^) littermate mice, 5 µg of protein were loaded per lane of a 3-8% gradient Tris-Acetate gel and bands were detected by chemiluminescence. The scanned Western blot (top) and a protein stain of the nitrocellulose membrane (bottom, dotted rectangle indicates the region represented in the upper panel in C), and estimated Munc13-1 protein levels normalized to total measured protein are shown in D, n = 3 brains each. **(E)** Western blot analysis of Munc13-1 levels from E18 or P0 whole brain tissue homogenates from different Munc13-1 knockout mouse lines. Lane 1, wild type (Munc13-1^+/+^); lane 2, homozygote germline recombined allele of the new Munc13-1 floxed (Munc13-1 cKO^gl^) mice; lane 3, Munc13-1 constitutive knockout (Munc13-1^-/-^) published in (Augustin et al., 1999); lane 4, Munc13-1 constitutive knockout (Munc13-1^mNeo/mNeo^) published in (Rhee et al., 2002); lane 5, Munc13-1/Munc13-2 double knockout (Munc13-1^-/-^+2^-/-^) published in (Varoqueaux et al., 2002). Munc13-1 cKO^gl^ exhibit a severe reduction of Munc13-1 protein levels comparable to that in well-characterized constitutive Munc13-1 knockout lines. Note that the Munc13-1^-/-^ (Augustin et al., 1999) mice express a truncated Munc13-1 protein product (*) that is detected by antibodies that bind N-terminal epitopes as discussed in Fig. 1 of (Man et al., 2015). A long exposure (middle) reveals that the anti-Munc13 antibody detects a protein band with slightly lower molecular weight in Munc13-1 knockout samples, which corresponds to ubMunc13-2 (Munc13-2) and is absent in the Munc13-1^-/-^+2^-/-^ sample. Munc13-1 cKO^gl^ animals express very low levels of a Munc13-1 protein product, likely corresponding to a protein lacking both exon 21 and 22, for which transcripts could be detected in brains of Munc13-1 cKO^gl^ mice using reverse transcriptase (rt) PCR (see methods). **(F)** Estimation of Munc13-1 levels in P0 whole brain homogenates from Munc13-1 wild type and littermate Munc13-1 cKO^gl^ animals indicates that the residual Munc13-1 protein is reduced to at most a few % of wild type Munc13-1 levels, n = 3 mice each. Data in D and F are mean ± SEM.

**Figure S3.**
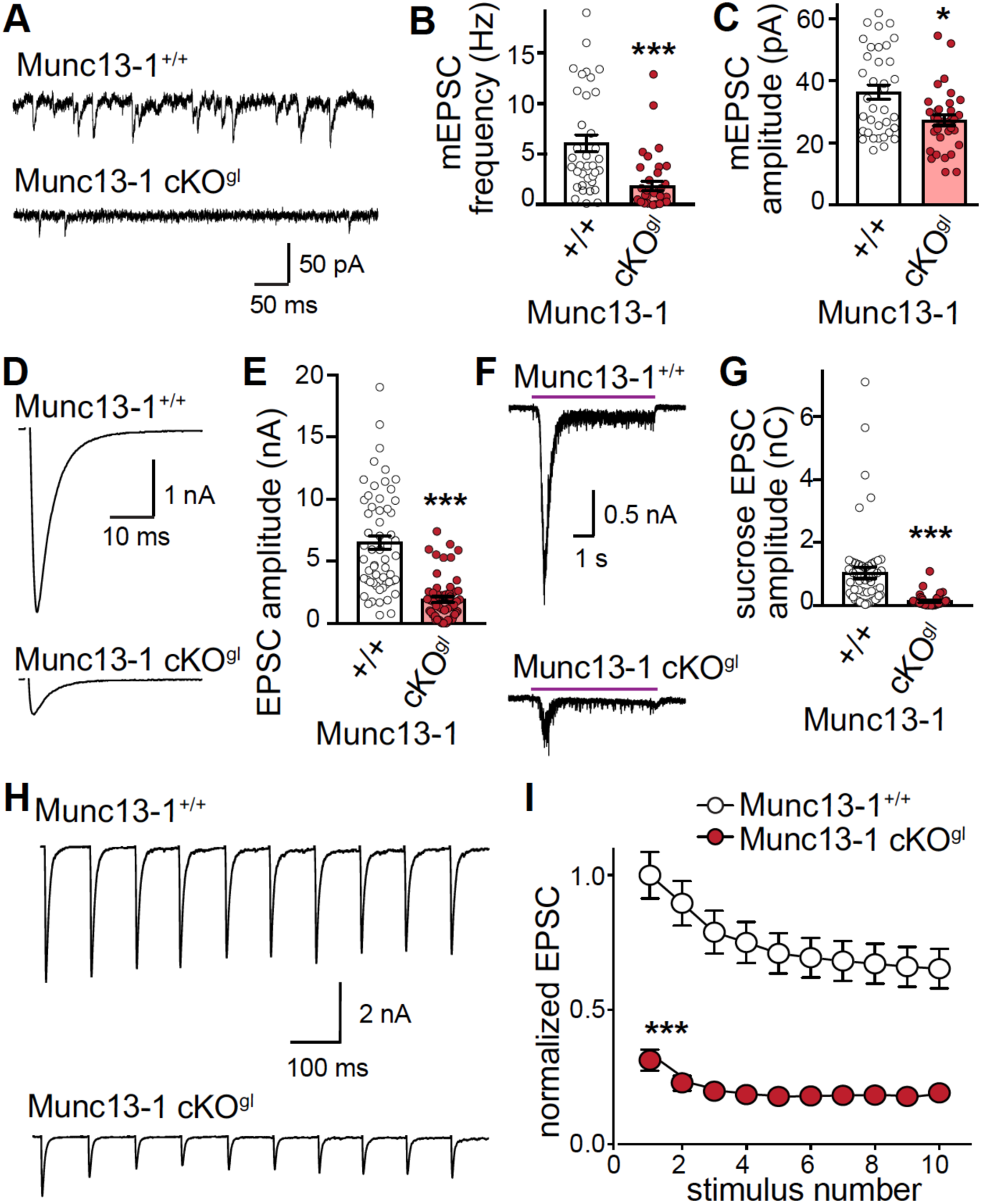
Munc13-1 deletion strongly impairs neurotransmitter release in autaptic hippocampal cultures, related to Fig. 3 **(A-C)** Sample traces (A) and quantification of miniature EPSC (mEPSC) frequency (B) and amplitude (C) recorded in presence of tetrodotoxin (TTX). B: Munc13-1^+/+^ = 36 neurons/3 independent cultures; Munc13-1 cKO^gl^ = 39/3; C: Munc13-1^+/+^ = 36/3; Munc13-1 cKO^gl^ = 33/3. **(D, E)** Sample traces (D) and quantification of average EPSC amplitudes (E) evoked by a 2 ms depolarization to 0 mV, Munc13-1^+/+^ = 57/3; Munc13-1 cKO^gl^ = 56/3. **(F, G)** Sample traces (F) and quantification of EPSCs (G) evoked by application of 500 mM sucrose. Munc13-1^+/+^ = 54/3; Munc13-1 cKO^gl^ = 53/3. **(H, I)** Sample traces (H) and quantification (I) of EPSCs triggered by 10 stimuli at 10 Hz. Quantification in I is shown normalized to the mean of the first EPSC of Munc13-1^+/+^, Munc13-1^+/+^ = 57/3; Munc13-1 cKO^gl^ = 56/3. Data are mean ± SEM, * p < 0.05, *** p < 0.001 as assessed by Mann Whitney test in B, C, E, G; two-way ANOVA (*** p < 0.001 for genotype and stimulus number) followed by Sidak’s multiple comparisons test *** p < 0.001 for all stimuli in I.

**Figure S4.**
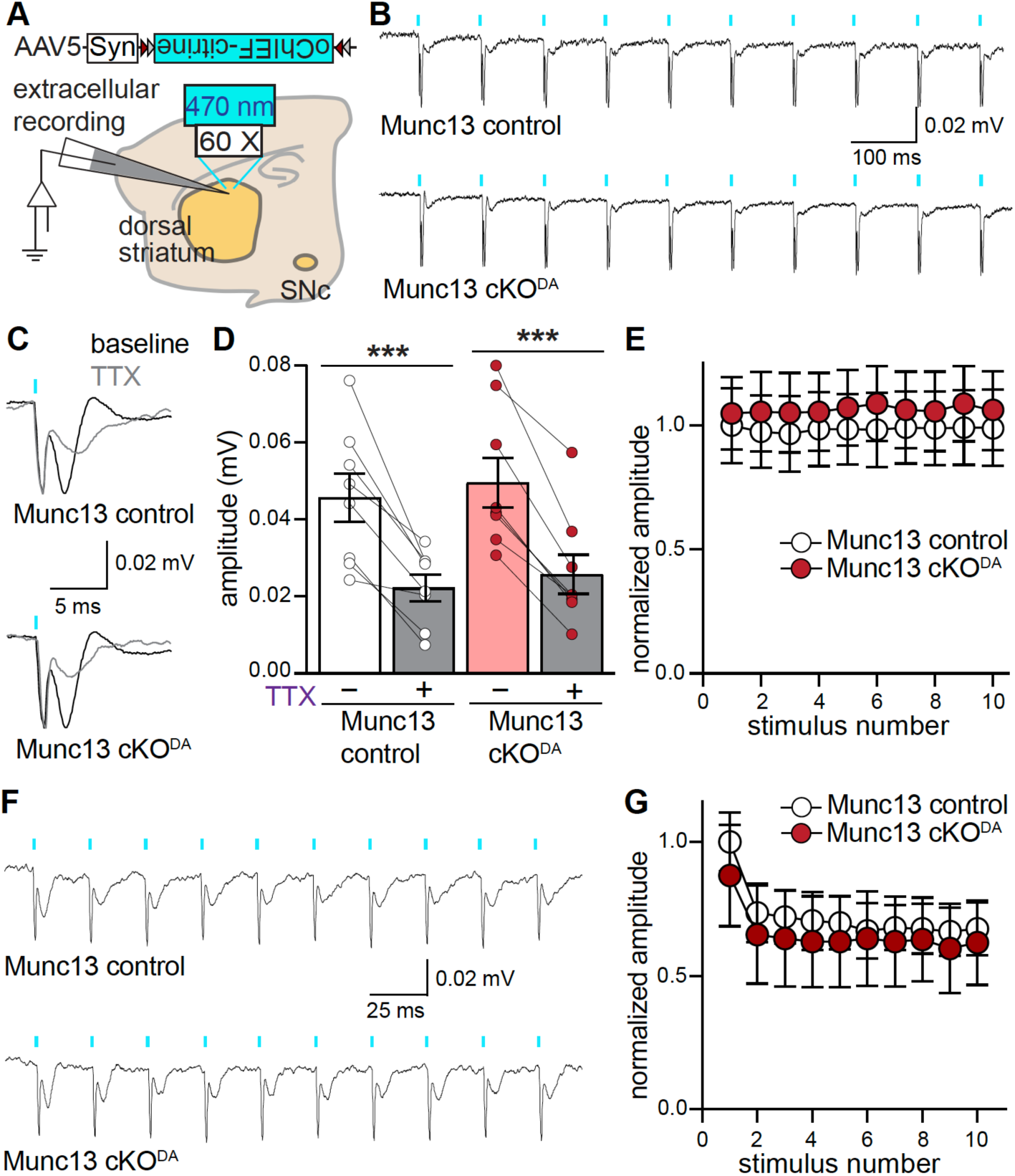
Munc13 cKO^DA^ does not impair dopamine axonal firing, related to Fig. 3 **(A)** Schematic outlining Cre-dependent expression of oChIEF for dopamine neuron activation and recording of extracellular field potentials in dorsolateral striatum. **(B-E)** Sample traces (B, C, average of 100 sweeps) and quantification (D, E) of extracellular potentials evoked by 10 stimuli at 10 Hz of optogenetic stimulation. In C, the extracellular potential for the first stimulus of a 10 Hz train before (black) and after 1 µM TTX (grey) is magnified. Quantification of extracellular potential amplitudes evoked by the 1^st^ stimulus of a 10 Hz train before and after TTX is shown in D, and the normalized extracellular potential amplitudes for 10 Hz after normalization to the mean first response amplitude of Munc13 control. The extent of TTX inhibition is similar between Munc13 control and cKO^DA^. Munc13 control = 8 slices/6 mice; Munc13 cKO^DA^ = 8/6. **(F, G)** Same as B and E but for the first 10 stimuli of a 40 Hz train, Munc13 control = 6/5; Munc13 cKO^DA^ = 6/5. Data are mean ± SEM, *** p < 0.001 as assessed by one-way ANOVA followed by Sidak’s multiple comparisons test in D; two-way ANOVA (p > 0.05 for genotype, stimulus number and interaction) in E; two-way ANOVA (*** p < 0.001 for stimulus number, p > 0.05 for genotype and interaction) followed by Sidak’s multiple comparisons test in G.

**Figure S5.**
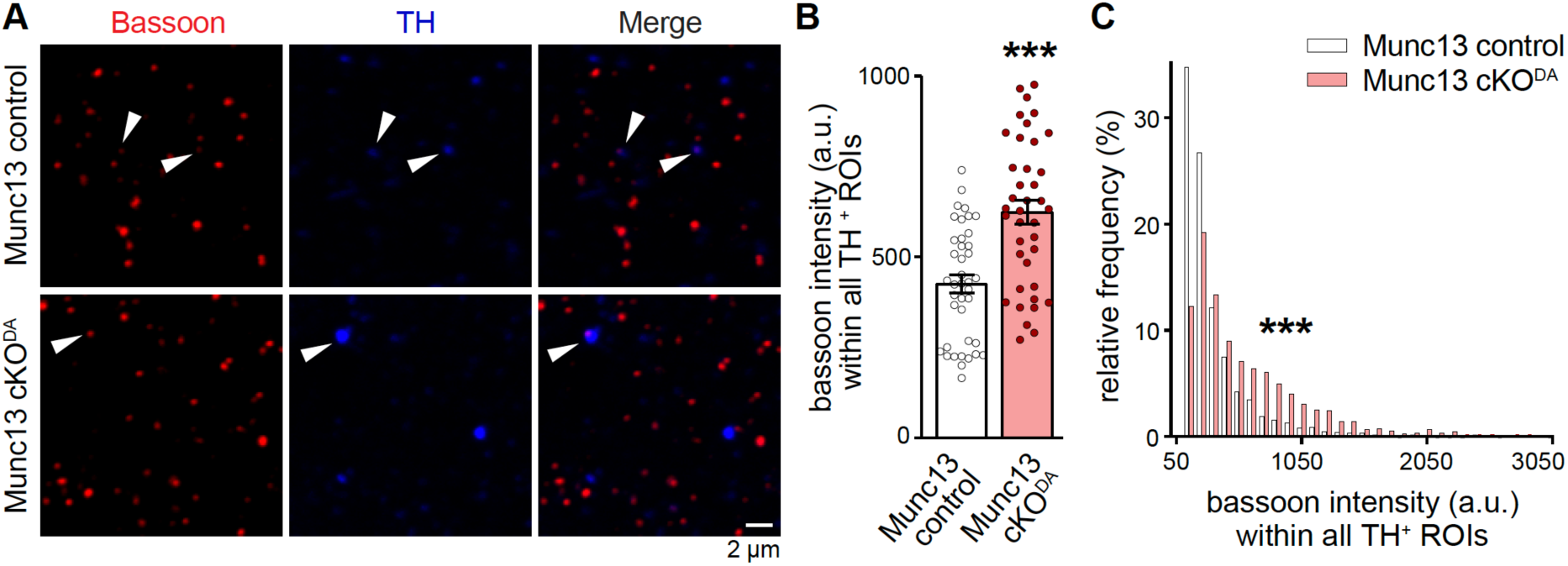
Bassoon clustering in dopamine synaptosomes, related to Fig. 5 **(A-C)** Sample confocal images (A) and quantification (B, C) of striatal synaptosomes stained with active zone marker bassoon (red) and dopamine axon marker TH (blue). Quantification of bassoon intensity within all TH ROIs (B) and frequency distribution histogram (C) are shown. Bassoon expression within TH positive synaptosomes is increased in Munc13 cKO^DA^. Only bassoon intensities within TH^+^ ROIs greater than 3 times the average intensity of all pixels are shown in B. The frequency histogram in C is plotted for all bassoon intensities within TH^+^ ROIs. Munc13 control = 40 images/4 mice; Munc13 cKO^DA^ = 39/4. Data in B are mean ± SEM, *** p < 0.001 as assessed by unpaired t-test in B and Kolmogorov-Smirnov test for data in C.

**Figure S6.**
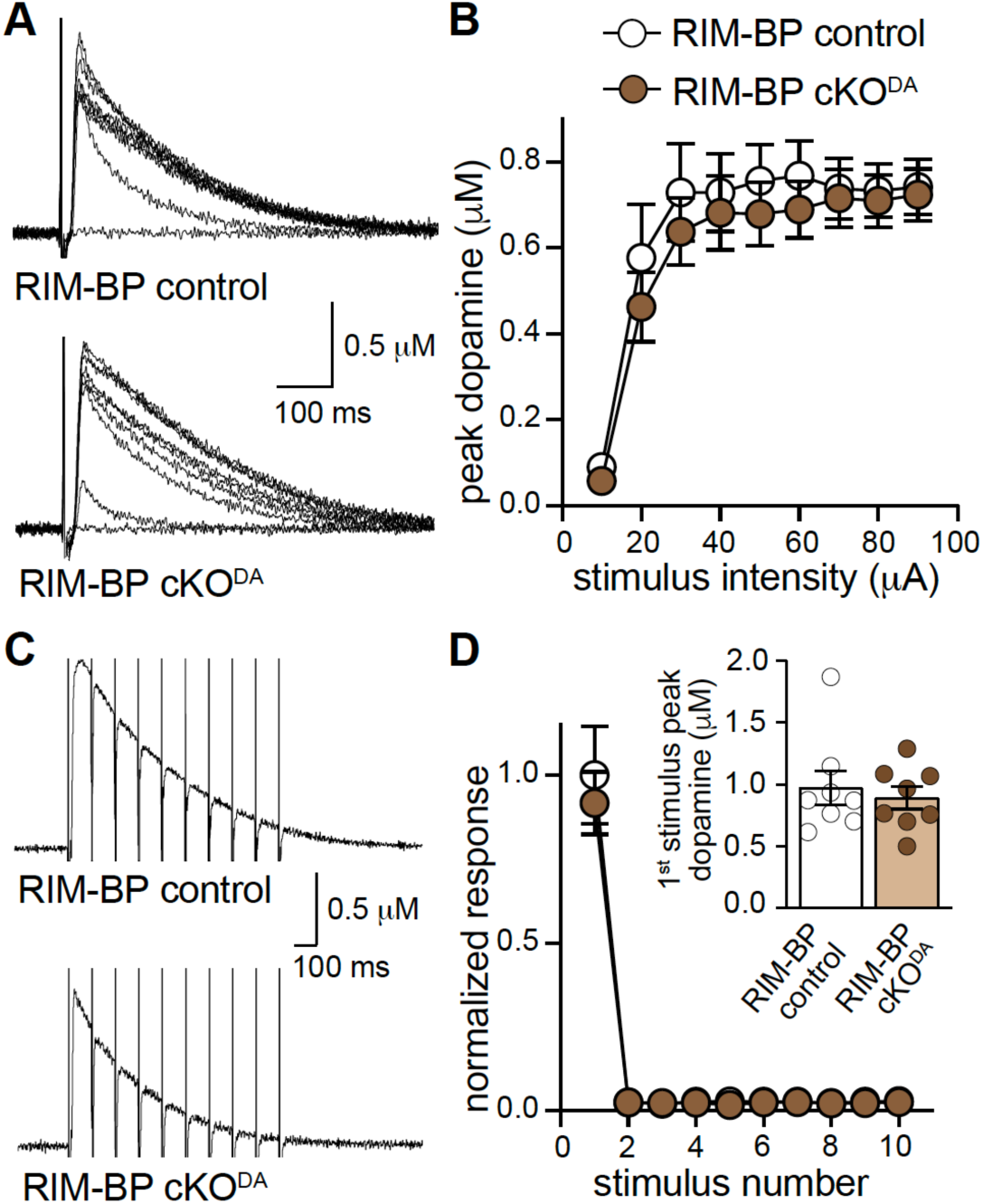
Deletion of RIM-BP1/2 does not impair electrically evoked dopamine release, related to Fig. 6 **(A, B)** Sample traces of (A, single sweeps) and quantification of peak amplitudes (B) of dopamine release evoked by electrical stimulation (10-90 µA single electrical pulses at increasing stimulation intensity), RIM-BP control = 16 slices/5 mice; RIM-BP cKO^DA^ = 16/5. **(C, D)** Sample traces (C, average of 4 sweeps) and quantification of peak amplitudes (D) of dopamine release normalized to the first peak amplitude of RIM-BP control in response to 10 stimuli at 10 Hz train, inset in D shows the peak dopamine amplitude for 1^st^ stimulus, RIM-BP control = 8/4; RIM-BP cKO^DA^ = 8/4. Data are mean ± SEM, p > 0.05 as assessed by two-way ANOVA (for genotype, stimulus and interaction) followed by Sidak’s multiple comparisons test in B, D and unpaired t-test for inset in D.

**Figure S7.**
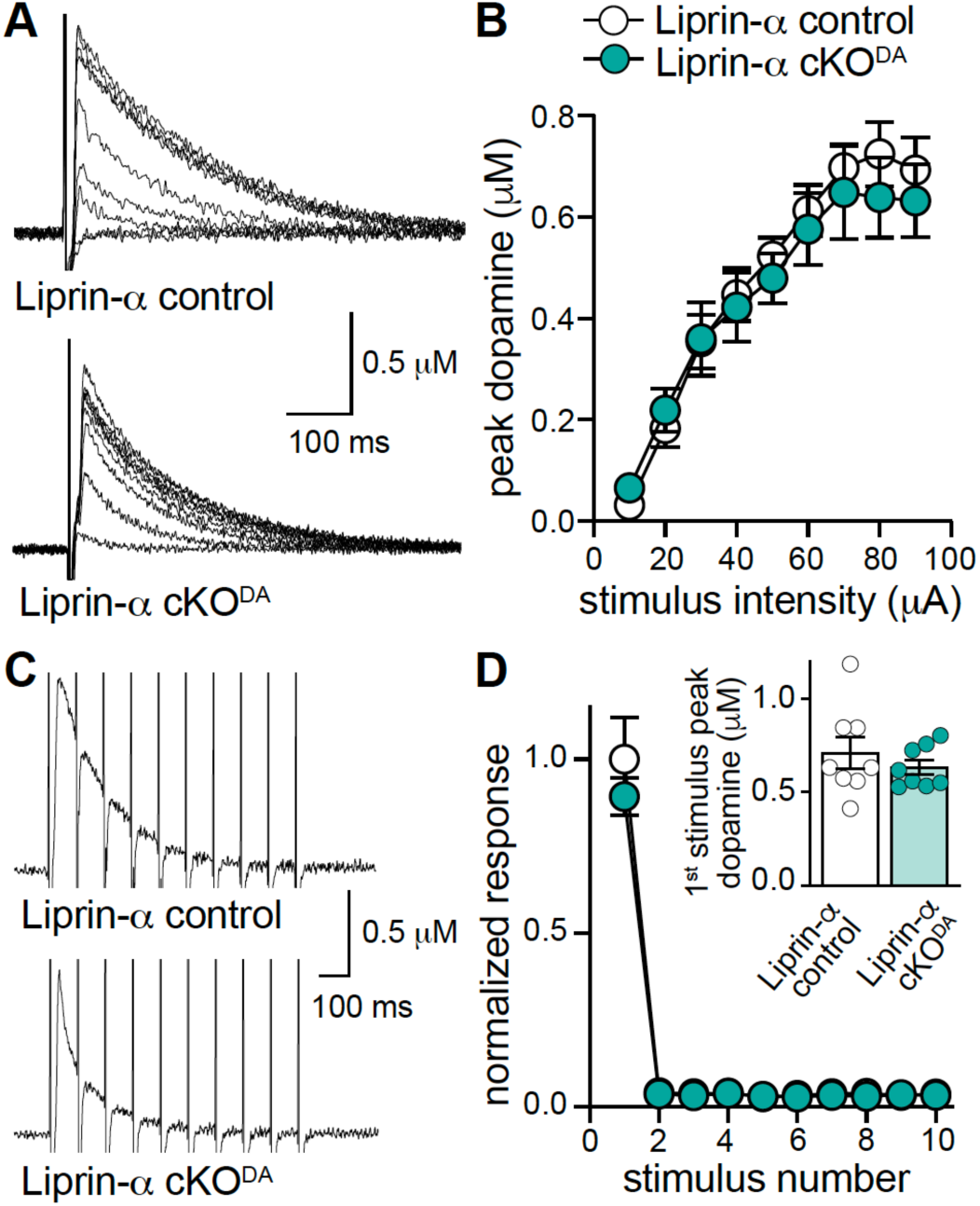
Deletion of Liprin-α2/3 does not impair electrically evoked dopamine release, related to Fig. 7. **(A, B)** Sample traces of (A, single sweeps) and quantification of peak amplitudes (B) of dopamine release evoked by electrical stimulation (10-90 µA single electrical pulses at increasing stimulation intensity), Liprin-α2/3 control = 12 slices/8 mice; Liprin-α2/3 cKO^DA^ = 12/8. **(C, D)** Sample traces (C, average of 4 sweeps) and quantification of peak amplitudes (D) of dopamine release normalized to the first peak amplitude of Liprin-α2/3 control in response to 10 stimuli at 10 Hz train, inset in D shows the peak dopamine amplitude for 1^st^ stimulus, Liprin-α2/3 control = 8/3; Liprin-α2/3 cKO^DA^ = 8/3. Data are mean ± SEM, p > 0.05 as assessed by two-way ANOVA (for genotype, stimulus and interaction) followed by Sidak’s multiple comparisons test in B, D and unpaired t-test for inset in D.

**Figure S8.**
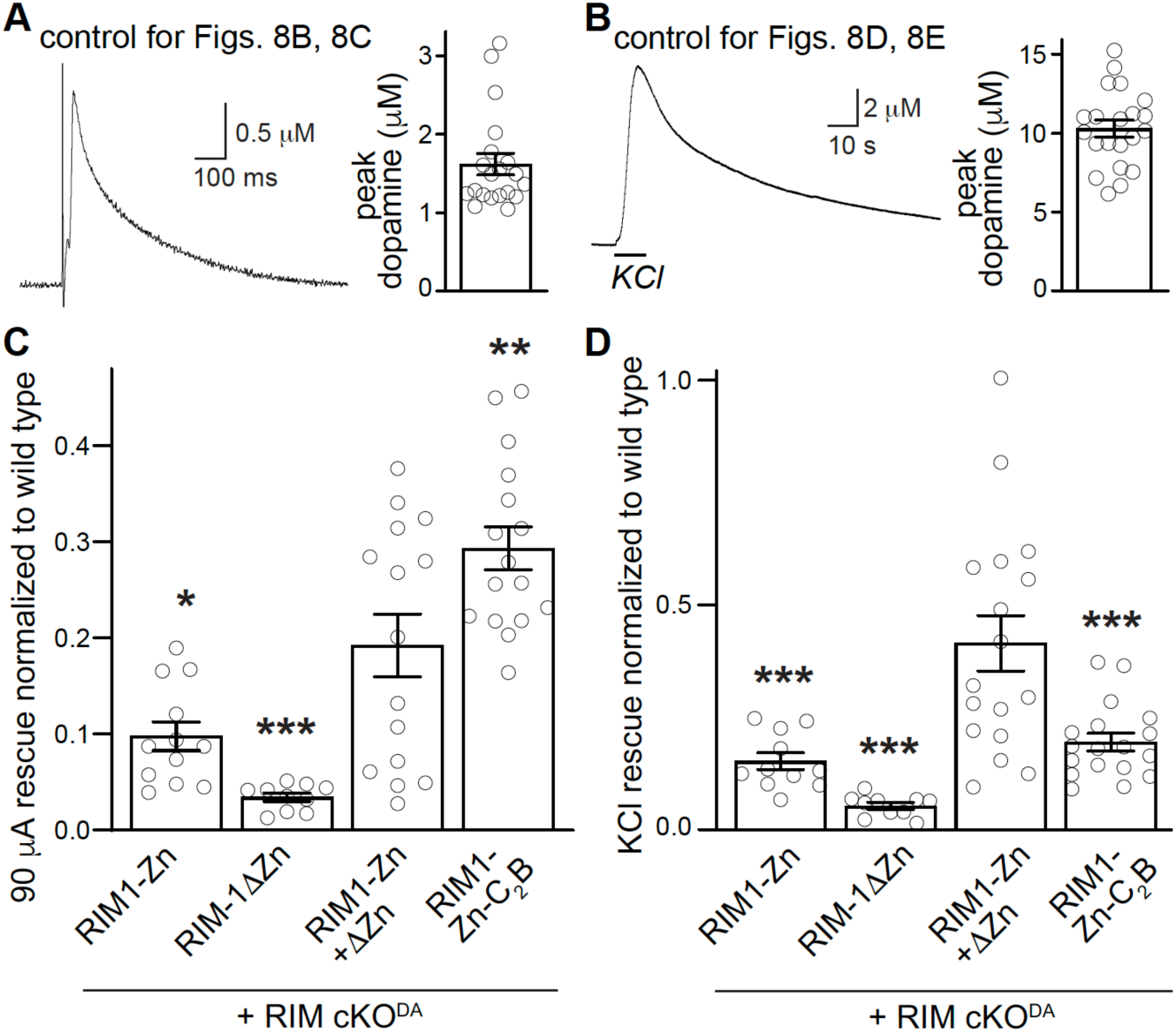
Control recordings and expression analyses for RIM1-Zn-HA-C_2_B rescue, related to Fig. 8 **(A)** Sample traces (single sweeps) and quantification of dopamine release evoked by a 90 µA electrical stimulus in slices of dorsolateral striatum in unrelated control animals recorded on the same days as the experiment shown in Fig. 8, n = 20 slices/4 mice. **(B)** Same as B but for a local 100 mM puff of KCl, n = 21/4. **(C)** Rescue of peak dopamine evoked by 90 µA electrical stimulus normalized to the average of wild type for all rescue conditions. RIM cKO^DA^ + RIM1-Zn = 12/4 mice; RIM cKO^DA^ + RIM1-ΔZn = 10/3; RIM cKO^DA^ + RIM1-Zn + RIM1-ΔZn = 15/4; RIM cKO^DA^ + RIM1-Zn-C_2_B = 16/4; wild type = 56/15 (used for normalization). **(D)** Same as D but for rescue estimated by KCl evoked dopamine release. RIM cKO^DA^ + RIM1-Zn = 11/4; RIM cKO^DA^ + RIM1-ΔZn = 10/3; RIM cKO^DA^ + RIM1-Zn + RIM1-ΔZn = 17/4; RIM cKO^DA^ + RIM1-Zn-C_2_B = 18/4; wild type = 55/15 (used for normalization). Data are mean ± SEM. * p < 0.05, ** p < 0.01, *** p < 0.001 as assessed by ANOVA followed by Dunnett’s multiple comparisons test in C and D where all comparisons were made to RIM cKO^DA^ + RIM1-Zn + RIM1-ΔZn.

## Notes

### Competing Interest Statement

The authors have declared no competing interest.

